# Mechanism underlying autoinducer recognition in the *Vibrio cholerae* DPO-VqmA quorum-sensing pathway

**DOI:** 10.1101/2019.12.19.881847

**Authors:** Xiuliang Huang, Olivia P. Duddy, Justin E. Silpe, Jon E. Paczkowski, Jianping Cong, Brad R. Henke, Bonnie L. Bassler

## Abstract

Quorum sensing is a bacterial communication process whereby bacteria produce, release and detect the accumulation of extracellular signaling molecules called autoinducers to coordinate collective behaviors. In *Vibrio cholerae*, the quorum-sensing autoinducer, DPO (3,5-dimethyl-pyrazin-2-ol), binds the receptor-transcription factor, VqmA. In response, the DPO-VqmA complex activates transcription of the *vqmR* gene encoding the VqmR small RNA. VqmR represses genes required for biofilm formation and virulence factor production. Here, we show that VqmA has DPO-dependent and DPO-independent activity. We solved the DPO-VqmA crystal structure and compared it to existing structures to understand the conformational changes the protein undergoes upon DNA binding. Analysis of DPO analogs reveals that a hydroxyl or carbonyl group at the 2’ position is critical for binding. The proposed DPO precursor, a linear molecule, Ala-AA (N-alanyl-aminoacetone), also binds and activates VqmA. DPO and Ala-AA occupy the same binding site as judged by site-directed mutagenesis and competitive ligand binding analyses.

---

Bacteria communicate and orchestrate collective behaviors using a process called quorum sensing (QS). QS relies on the production, release, detection, and group-wide response to extracellular signaling molecules called autoinducers (AIs) that accumulate with increasing cell density (reviewed in 1). In the global pathogen *Vibrio cholerae*, multiple QS pathways converge to repress biofilm formation and virulence factor production at high cell density (HCD) (1–6). One of the *V. cholerae* QS pathways is composed of an AI, 3,5-dimethyl-pyrazin-2-ol (DPO), and its partner cytoplasmic receptor, VqmA. VqmA is a transcription factor that, upon binding DPO, activates expression of a gene encoding a small RNA called VqmR (7,8). VqmR post-transcriptionally controls many target genes (7,9). Of note, VqmR represses translation of *ctx* and *rtxA*, two toxin-encoding genes, *vpsT*, encoding a component required for biofilm formation, and *aphA*, encoding the low cell density (LCD) QS master regulator AphA (7,9). Thus, analogous to the previously discovered *V. cholerae* QS systems, the consequence of DPO binding to VqmA at HCD is repression of virulence and biofilm formation.

Homologs of the VqmA receptor are restricted to the *Vibrio* genus, and the vibriophage VP882 (10). By contrast, threonine dehydrogenase (Tdh), an enzyme required for DPO biosynthesis, is highly conserved among bacteria, archaea, and eukarya (11). Consistent with this distribution, with respect to bacteria, DPO is produced by multiple species in different orders (8). Tdh oxidizes L-threonine to 2-amino-3-ketobutyric acid (AKB), which spontaneously decarboxylates to aminoacetone (12). In the proposed mechanism for DPO biosynthesis, aminoacetone is predicted to undergo a condensation reaction with L-alanine to yield N-alanyl-aminoacetone (Ala-AA) (8). This putative linear precursor is subsequently converted to DPO by an unknown mechanism.

In this report, we characterize ligand-controlled activation of VqmA in *V. cholerae*. In contrast to canonical LuxR-type QS receptor-transcription factors, which typically require binding to their cognate ligands to fold and activate transcription (13–16), here, we find that VqmA does not require DPO for folding or basal transcriptional activity, but does require binding DPO to drive maximal activity. These same properties are relevant for the phage VP882-encoded VqmA homolog (VqmAPhage), suggesting that ligand-independent activity is a conserved feature among VqmA proteins. We show that VqmA binds to and is activated by both DPO and the putative DPO intermediate, Ala-AA. We solved the crystal structure of VqmA bound to DPO. Comparison to existing structures (17,18) allowed us to provide insight into the conformational changes VqmA undergoes upon DNA binding. We were unable to obtain the crystal structure of VqmA bound to Ala-AA because Ala-AA spontaneously cyclizes into DPO under our crystallization conditions. Mutagenesis of the VqmA ligand-binding pocket shows that DPO and Ala-AA likely occupy the same site in VqmA, as substitutions in residues that alter DPO binding and activity have similar effects on Ala-AA binding and activity. Characterization of binding by isothermal titration calorimetry (ITC)-based competition experiments and kinetic studies reveals that VqmA prefers DPO over Ala-AA. Assessment of the activities of a panel of DPO analogs revealed that a 2’ hydroxyl or carbonyl group is required for binding to VqmA. Taken together, our results show that VqmA is promiscuous with respect to DPO and Ala-AA. Nonetheless, maximal VqmA activity depends on ligand binding. The ability of VqmA to activate gene expression to different levels in the unbound state and when bound to two different ligands could enable greater flexibility in regulating gene expression than exists in canonical LuxR-type QS receptors, which typically have no activity in the Apo-state and show strict specificity for a single partner ligand.

## RESULTS

### VqmA possesses both ligand-independent and ligand-dependent activity

VqmA is a cytoplasmic transcription factor and QS receptor that binds the AI DPO, a hydroxypyrazine that requires the Tdh enzyme for its production (8). The VqmA receptor is restricted to vibrios and one vibriophage. By contrast, canonical LuxR-type cytoplasmic QS receptor-transcription factors exist in thousands of Gram-negative bacterial species and they bind AIs that are acyl-homoserine lactones (AHLs) (reviewed in 1). AHL AIs are produced by LuxI-type AI synthases. LuxR-type receptors typically require binding to their cognate AIs to properly fold, become soluble, and activate transcription (13–16). Thus, LuxI-type AI synthases and LuxR-type receptors function as obligate pairs, and mutation of the *luxI* gene confers the identical defect as does mutation of the *luxR* gene (19). We wondered whether the Tdh-VqmA pair functions in an analogous manner. To probe this question, we constructed a reporter system to monitor activation of transcription by VqmA. We fused the *lux* (luciferase) operon to the *vqmR* promoter (P*vqmR-lux*) and integrated it onto the *V. cholerae* chromosome at an ectopic locus (*lacZ*). We measured transcriptional activity in different *V. cholerae* strains and across different conditions. WT *V. cholerae* harboring the P*vqmR-lux* reporter exhibited maximal activity at HCD, which is consistent with our previous report (Fig. 1A and (8)). The Δ*tdh* mutant, which is defective for DPO production, elicited ~10-fold lower activity (Fig. 1A). Restoration of light production to approximately the WT level occurred in the Δ*tdh* mutant following exogenous addition of 1 μM DPO (Fig. 1A). In the Δ*vqmA* strain, P*vqmR-lux* levels were ~330-fold lower than in the WT and ~23-fold lower than in the Δ*tdh* strain to which no DPO was added (Fig. 1A). Exogenous addition of DPO did not induce light production in the Δ*vqmA* strain (Fig. 1A). Thus, unlike what is observed for canonical LuxI-LuxR-type systems, the Δ*tdh* synthase mutant and the Δ*vqmA* receptor mutant have different phenotypes with respect to QS output.

**Figure 1.**
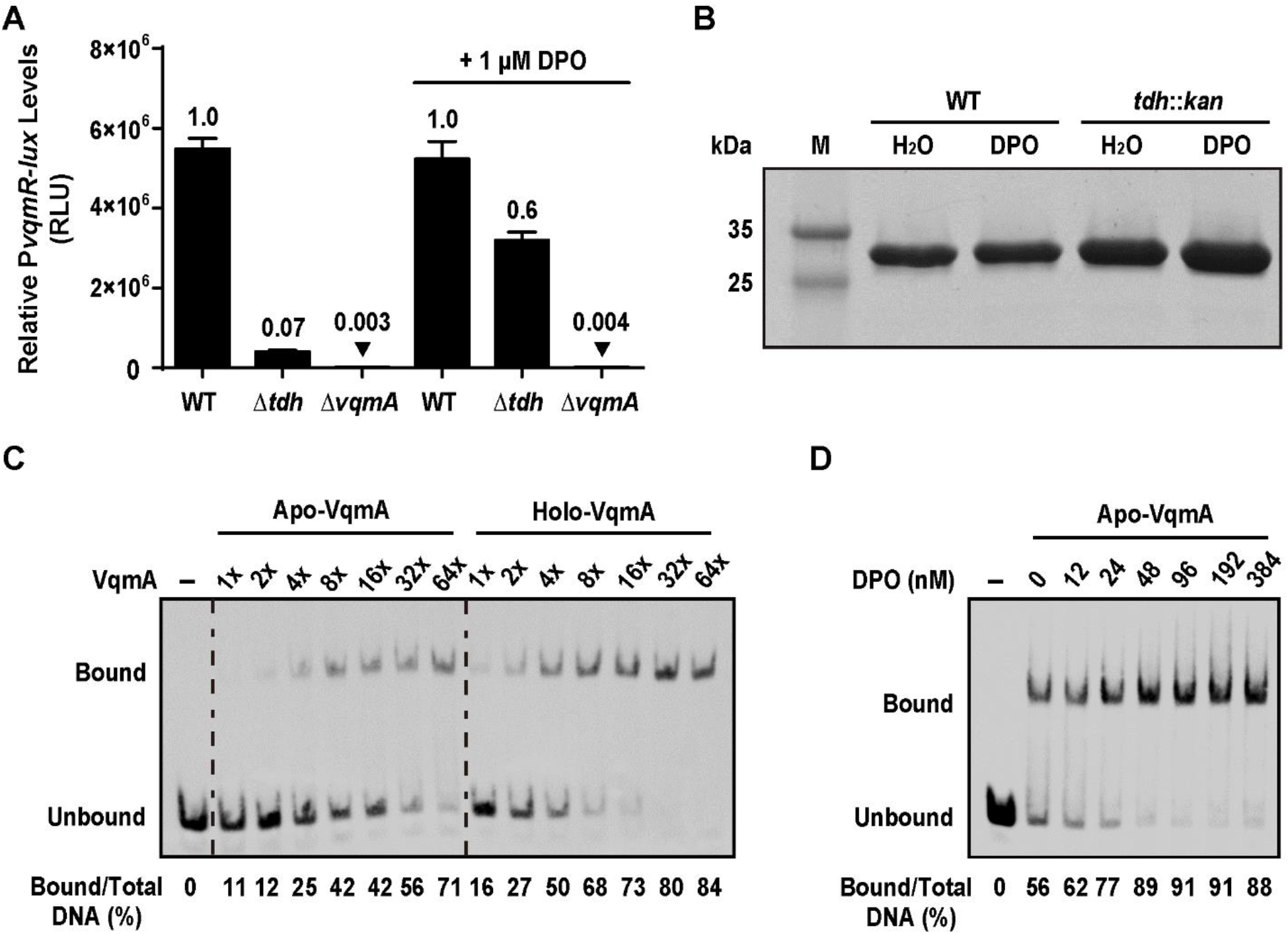
VqmA is active in the absence of DPO. (A) P*vqmR-lux* reporter activity for the indicated *V. cholerae* strains after 6 h of growth in the absence or presence of 1 μM DPO. Data represented as mean ± SD with n = 3 biological replicates. (B) SDS-PAGE gel of purified His-VqmA protein produced in WT or *tdh*::*kan E. coli* without or with 100 μM DPO added during expression. M = molecular weight marker. (C) EMSA showing VqmA binding to P*vqmR* promoter DNA. 540 pM biotinylated P*vqmR* DNA and either no protein (Lane 1, designated -), Apo-VqmA (Lanes 2-8), or Holo-VqmA (Lanes 9-15). Relative protein concentrations are indicated: 1X = 2 nM and 64X = 130 nM. (D) EMSA showing binding of 50 nM Apo-VqmA (lanes 2-8) to P*vqmR* DNA in the absence and presence of the indicated amounts of DPO. Lane 1 as in (C). In panels (C) and (D), quantitation of the percent P*vqmR* DNA bound appears below each lane.

Two possibilities could underpin the differences in the *lux* phenotypes of the Δ*vqmA* and Δ*tdh* mutants: either DPO is produced via a Tdh-independent mechanism, or VqmA displays basal activity in the absence of its ligand. To investigate the first possibility, we quantified DPO production by WT, Δ*tdh*, and Δ*vqmA V. cholerae*. Cell-free culture fluids prepared from each strain were supplied to the *V. cholerae* Δ*tdh* P*vqmR-lux* reporter strain and light production was measured (Fig. S1A). Activity present in the preparations was compared to that following addition of known amounts of synthetic DPO. The EC_50_ for synthetic DPO is ~1 μM (8). Using this value, we could estimate that the WT and Δ*vqmA* culture fluids contained ~1-2 μM DPO while the amount of DPO in culture fluid from the Δ*tdh* strain was below the 60 nM detection limit of the reporter (Fig. S1A). Thus, if any DPO is produced by a Tdh-independent mechanism(s), it is insufficient to elicit the order-of-magnitude difference in P*vqmR* activity between the Δ*vqmA* and Δ*tdh V. cholerae* strains in Fig. 1A. To test the second possibility, that VqmA is stable and possesses measurable activity in the absence of DPO, we produced VqmA in WT (i.e., *tdh*^+^) and *tdh::kan E. coli* and assessed its solubility. *E. coli*, like *V. cholerae*, produces DPO in a Tdh-dependent manner (8). Fig. 1B shows that comparable amounts of soluble VqmA were obtained from the WT and *tdh*::*kan* strains, irrespective of the addition of DPO. To eliminate the possibility that a trace amount of DPO is produced by the *tdh*::*kan* strain, or that some other ligand stabilizes VqmA in *tdh::kan E. coli*,we produced hexahistidine-tagged VqmA (His-VqmA) via *in vitro* translation using a cell-free system. *In vitro* translated His-VqmA protein was readily produced and soluble (Fig. S1B). Thus, we conclude that VqmA does not depend on a ligand to properly fold.

In *V. cholerae*, VqmA is produced constitutively whereas *vqmR* is expressed predominantly at HCD (8). Given our finding that the solubility of VqmA is unaffected by DPO (Fig. 1B), we hypothesized that the VqmA DNA-binding capability could be enhanced by DPO. To examine VqmA DNA-binding as a function of ligand occupancy, we purified VqmA from *tdh*::*kan E. coli* that had been grown in the presence or absence of exogenously supplied DPO (100 μM). We assessed Apo- and Holo-VqmA binding to P*vqmR* DNA using electromobility shift assays (EMSAs). Our results show that, at sufficient protein concentrations, Apo-VqmA shifted P*vqmR* DNA (Fig. 1C), consistent with our finding that VqmA possesses basal activation of P*vqmR* in the Apo-state. We found that 2-8-fold more protein was required to achieve a ~50% shift of the P*vqmR* DNA with Apo-VqmA compared to Holo-VqmA (Fig. 1C). Thus, Holo-VqmA has a higher affinity for P*vqmR* than Apo-VqmA. Furthermore, addition of DPO to Apo-VqmA increased DNA binding to P*vqmR* in a DPO concentration-dependent manner (Fig. 1D). We conclude that the basal DNA-binding activity of Apo-VqmA is enhanced when VqmA is bound to DPO. These results explain why the *V. cholerae* Δ*tdh* mutant exhibits P*vqmR-lux* expression while the Δ*vqmA* mutant does not (Fig. 1A).

Considering the differences in the biochemical properties of the VqmA receptor relative to LuxR-type QS receptors, we wanted to examine whether their control of QS outputs also differed. To do this, we measured the effect of DPO binding to VqmA on P*vqmR* activation, representing the *V. cholerae* VqmA pathway, and compared the results to the well characterized LasI-LasR QS system from *Pseudomonas aeruginosa*. Briefly, in *P. aeruginosa*, upon binding 3OC_12_-homoserine lactone (3OC_12_-HSL), LasR activates transcription of many genes including, germane to our studies, *lasB* (20). Here, we introduced the relevant *P. aeruginosa* and *V. cholerae* QS components into *E. coli* (Fig. 2). Specifically, we used Δ*tdh E. coli* as the host for both reporter systems to eliminate native AI production (*E. coli* does not produce 3OC_12_-HSL). Into this strain, we introduced a pair of plasmids encoding either arabinose-inducible *vqmA* and P*vqmR-lux*, or arabinose-inducible *lasR* and P*lasB-lux*. Consistent with our finding in Δ*tdh V. cholerae* (Fig. 1A), *E. coli* producing VqmA in the absence of DPO generated 15-fold more P*vqmR* reporter activity than a strain lacking the vector harboring arabinose-inducible *vqmA* (Fig. 2). In response to exogenous DPO, P*vqmR*-driven light production increased an additional 3-fold. LasR, by contrast, expressed minimal P*lasB-lux* production in the absence of 3OC_12_-HSL relative to the no-vector control. In response to 3OC_12_-HSL, a 143-fold increase in P*lasB-lux* expression occurred (Fig. 2). Taken together, these results show that both VqmA and LasR are maximally activated by their respective AIs. However, VqmA activity, in contrast to LasR activity, is not strictly dependent on an AI. In the opposite vein, the relative activation achieved by 3OC_12_-HSL-LasR is almost two-orders of magnitude greater than that for the DPO-VqmA complex.

**Figure 2.**
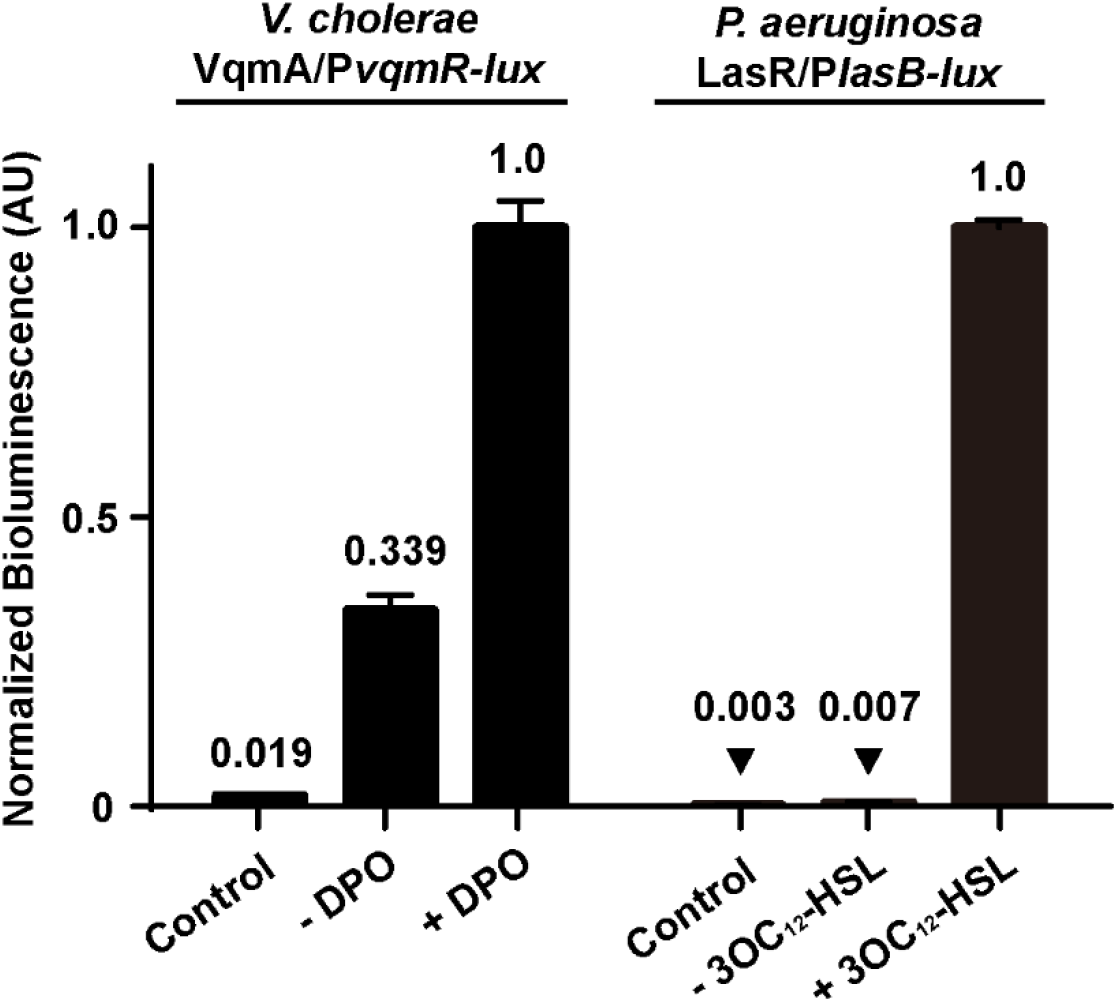
VqmA exhibits activity in the absence of its cognate AI and the LasR QS receptor does not. Normalized reporter activity from *E. coli* harboring the VqmA/P*vqmR-lux* or the LasR/P*lasB-lux* system. Left: *E. coli* carrying one plasmid with P*vqmR-lux* and a second plasmid with pBAD-*vqmA*. The strain in the lane designated control lacks the pBAD-*vqmA* plasmid. Right: *E. coli* carrying one plasmid with P*lasB-lux* and a second plasmid with pBAD-*lasR*. The strain in the lane designated control lacks the pBAD-*lasR* plasmid. All strains were grown in the presence of 0.075% arabinose and in the absence or presence of the designated AI (20 μM DPO or 1 μM 3OC_12_-HSL). AIs were added to the control strains. Data represented as mean ± SD with n = 3 biological replicates.

We wondered if the ligand-independent response we observed for *V. cholerae* VqmA extended to other VqmA proteins, such as that from the phage VP882-encoded VqmA homolog, VqmA_Phage_. VqmA_Phage_ binds host-produced DPO and, in response, activates expression of the gene, *qtip*. Qtip promotes host cell lysis (10). Similar to *V. cholerae* VqmA, VqmA_Phage_ activated P*qtip* transcription in the absence of DPO, and activation was enhanced approximately 3-fold when DPO was added (Fig. S1C). Thus, the VqmA AI-independent function extends to the phage-encoded system and is likely an inherent property of VqmA proteins more generally.

### The proposed linear DPO precursor Ala-AA binds to and activates VqmA

As noted, Tdh is required to convert L-threonine to AKB, which spontaneously decarboxylates to aminoacetone. Aminoacetone is presumed to condense with L-alanine to form the putative DPO precursor Ala-AA (8) (Fig. S2A). Consistent with the placement of threonine upstream and Ala-AA downstream of Tdh in this pathway, we previously showed that exogenous addition of Ala-AA, but not threonine, restores P*vqmR-lux* expression in Δ*tdh V. cholerae* to WT levels (8). This finding suggested one of two possibilities: first, in our cell-based assay, the cells convert Ala-AA to DPO, and newly-produced DPO drives P*vqmR-lux* expression. Alternatively, Ala-AA could bind to VqmA and induce P*vqmR-lux* expression without conversion to DPO. To distinguish between these possibilities, we performed EMSAs using Apo-VqmA to which we added either threonine, DPO, or Ala-AA (Fig. 3A). As anticipated, provision of L-threonine did not increase VqmA binding relative to when water was added (Fig. 3A). By contrast, inclusion of DPO increased the percentage of P*vqmR* DNA bound to VqmA ~2-5 fold (Fig. 3A). Surprisingly, the consequence of adding Ala-AA was indistinguishable from that of DPO (Fig. 3A). That is, Ala-AA also drives VqmA to bind P*vqmR. In vitro* translated His-VqmA behaved identically with respect to the ligands (Fig. S2B). Importantly, we excluded the possibility that Ala-AA is converted to DPO under our EMSA conditions using MS (DPO: 125 *m/z* and Ala-AA: 145 *m/z*). In each case, the ligand added, DPO or Ala-AA, was the only ligand detected by MS (Fig. S2C). Taken together, we conclude that both DPO and Ala-AA bind and enhance VqmA DNA binding and transcriptional activity.

**Figure 3.**
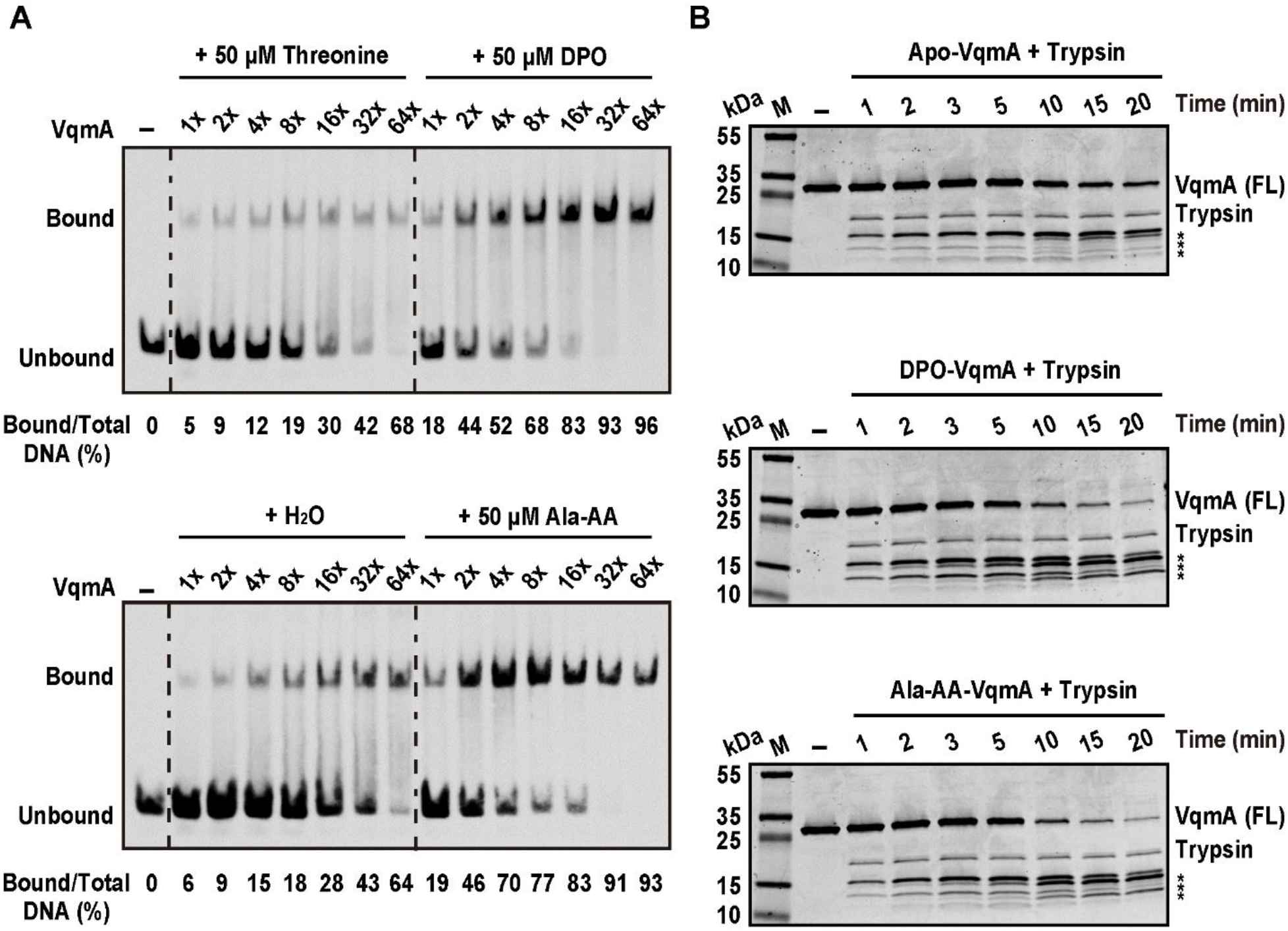
Ala-AA, the proposed linear precursor of DPO, binds to and activates VqmA. (A) EMSA of VqmA binding to P*vqmR* DNA in the presence of threonine, DPO, water, or Ala-AA. Probe and protein concentrations as in Fig. 1 (C). (B) Trypsin digestion of VqmA in the absence (H_2_O, top) or presence of ligand (DPO, middle; Ala-AA, bottom). M = molecular weight marker, FL = full-length. Time points (min) are indicated above each lane. FL VqmA (~27 kDa) and trypsin (~23 kDa) are marked. Asterisks denote VqmA proteolytic cleavage products.

To explore whether binding of DPO and Ala-AA to VqmA drives the same or different protein conformational changes, we performed limited proteolysis. Apo-, DPO-bound, and Ala-AA-bound VqmA were all digested by trypsin. However, the DPO-bound and Ala-AA-bound VqmA proteins were more susceptible to trypsin digestion than Apo-VqmA (Fig. 3B). These results indicate that a conformational change occurs upon ligand binding, and, moreover, that the change is similar for both ligands. Indeed, in companion experiments, we assessed the binding of DPO and Ala-AA to Apo-VqmA using thermal shift analysis. Both ligands caused similar increases (about 6°C) in thermal stability of VqmA (Fig. S2D). Our data support a model in which both DPO and the linear Ala-AA molecule bind VqmA and effectively activate its transcriptional activity.

The above results show that Ala-AA and DPO have identical activities in both our *in vivo* cell-based and *in vitro* protein-based assays. One caution, however, is that Ala-AA is the presumed DPO-precursor, but its *in vivo* role has not been verified. After DPO was discovered and threonine and alanine were shown to be incorporated into DPO, the prediction of Ala-AA as the DPO-precursor was put forward based on common chemical reactions in which amino acids and amino-containing compounds participate (8). Here, we aimed to prove or disprove whether Ala-AA is indeed the biological precursor to DPO. To do this, we conducted isotopic labeling experiments to determine whether Ala-AA cyclizes into the final DPO product. We administered deuterium-labeled Ala-AA (d_3_-Ala-AA) ((*S*)-2-amino-N-(2-oxopropyl)-3,3,3-d_3_-propanamide, shown in Fig. S3A) to Δ*tdh V. cholerae* and analyzed incorporation of the deuterium label into DPO by LC-MS. As controls, we also administered d_3_-Ala-AA to: (1) medium alone to eliminate the possibility of spontaneous conversion in the absence of cells; (2) Δ*tdh* Δ*vqmA V. cholerae* to confirm that binding to VqmA is not required for conversion; and (3) to WT *V. cholerae* to show that the amount of labeled DPO (d_3_-DPO) extracted is identical between the Tdh^+^ and Tdh^-^ strains to verify that Tdh is not functioning in any additional downstream step in the pathway. We detected d_3_-DPO in extracts at approximately 5% of the total concentration of added d_3_-Ala-AA (Fig. S3B). However, analysis of our synthetic preparations of Ala-AA and d_3_-Ala-AA showed that approximately 5% DPO and d_3_-DPO, respectively, are present (Fig. S3C) as irremovable impurities. The presence of small amounts of DPO demonstrated the facile nature of Ala-AA to undergo cyclization at neutral pH or basic conditions. Thus, in the labeling experiments, the low level of d_3_-DPO present in the extracts can be attributed to the starting material and it is not a consequence of enzymatic conversion of d_3_-Ala-AA to d_3_-DPO (Fig. S3C). There was no case in which administration of d_3_-Ala-AA to cells resulted in accumulation of recovered DPO with a 3 Da mass shift above the background level. This result suggests that, at least under our conditions, exogenously supplied Ala-AA is not converted into DPO *in vivo* (Fig. S3B). We suspect that some other compound could be the precursor to DPO and we are working to discover its identity. Nonetheless, our findings with Ala-AA are curious as they show that the VqmA receptor binds to and is activated by an apparently non-pathway substrate. We return to ideas concerning the ramifications of these findings in the Discussion.

### Structural and biochemical characterization of VqmA

To explore the molecular basis underlying VqmA binding to ligands, we solved the crystal structure of VqmA bound to DPO in the absence of DNA to 2.0 Å using multiwavelength anomalous diffraction (Table S1). The structure of the DPO-VqmA complex bound to DNA was reported recently (17). Additionally, during preparation of this manuscript, the structure of DPO-VqmA without DNA was reported with a similar conformation as in our structure (18). Together, the three structures enabled comparisons to discover the conformation changes VqmA undergoes upon DNA binding, and in the context of our work, to gain insight into how VqmA could bind both DPO and Ala-AA.

The DPO-VqmA complex is a head-to-head dimer with 1:1 protein:ligand stoichiometry. The structure revealed two distinct domains: an N-terminal PAS (Per-Arnt-Sim) ligand-binding domain (LBD) and a C-terminal HTH (helix-turn-helix) DNA-binding domain (DBD) (Fig. 4A). The N-terminal PAS LBD consists of a characteristic five-stranded antiparallel β-sheet core, which forms the DPO binding pocket along with several α-helices and loops, and this domain contains the bulk of the buried surface area of the dimerization interface (Fig. 4B, C, respectively). The PAS domain is connected to the C-terminal HTH DBD by a long flexible loop region. The DBD has a stereotypical HTH tetra-helical fold, which is common in transcription factors, including LuxR-type QS receptors (Fig. 4D). Both the LBD PAS and DBD HTH domains are common across biology. However, according to the DALI server (21), which uses structures from the Protein Data Bank to assess structural homologies based on fold rather than amino acid sequence, protein structures containing a domain composition similar to VqmA have not otherwise been reported. The closest VqmA homologs based on fold either contain the PAS domain as part of a signal transduction protein (i.e. TodS and VraR) or the HTH domain as part of a transcription factor (i.e. CviR and VpsT), but not both (Table S2).

**Figure 4.**
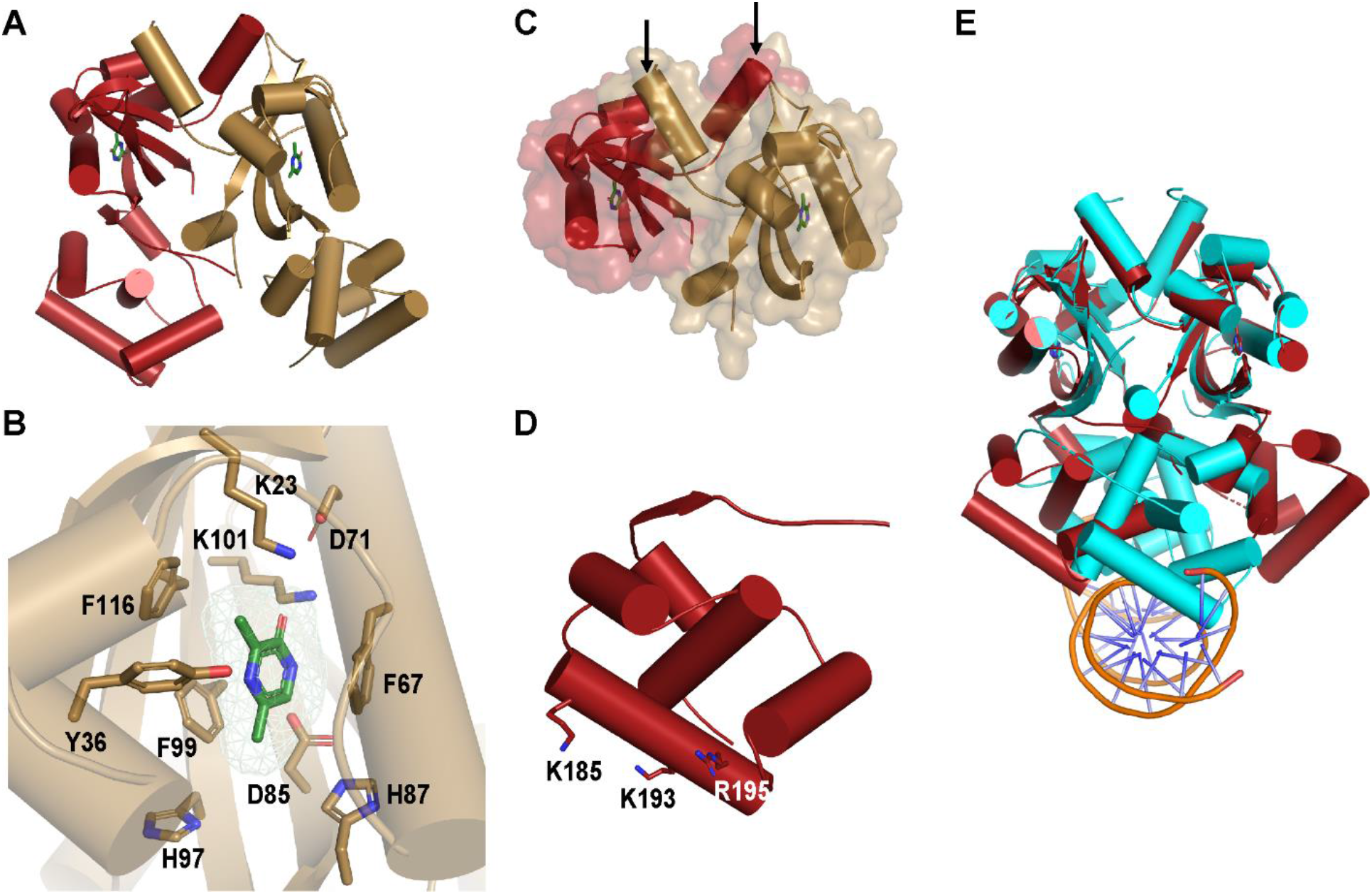
Crystal structure of VqmA bound to DPO. (A) 2.0 Å crystal structure of full-length VqmA as a dimer (monomers labeled in red and gold) with two molecules of DPO (green) bound. (B) Zoomed in view of one monomer (gold) showing the DPO ligand (green) and its spatial relationship to the residues tested in the mutagenesis studies (Table 1 and Fig. S6). The DPO-VqmA interface is primarily stabilized by residues F67, F99, and K101. (C) The N-terminal PAS LBD of VqmA in cartoon format as in (A) showing the buried surface area between monomers. The dimer is stabilized by extensive buried surface region and crossover of two helices (highlighted by the arrows) at the N-termini (residues 7-15) of the two monomers. (D) The C-terminal HTH DBD of VqmA in cartoon format as in (A). Based on solvent accessibility, K185, K193, and R195 are proposed to be involved in DNA binding. (E) Structural comparison of VqmA-DPO (red) with a previously solved structure of VqmA-DPO bound to DNA (PDB: 6IDE (17); protein: cyan; DNA: orange and blue).

**Table 1.** K_d_ values for WT and mutant VqmA proteins for DPO and Ala-AA.

| Mutation | DPO (μM) | Ala-AA (μM) |
| --- | --- | --- |
| WT | $2.3 \pm 0.1$ | $14.0 \pm 2.7$ |
| K23R | $1.2 \pm 0.6$ | $4.9 \pm 0.8$ |
| Y36F | $8.8 \pm 2.4$ | $6.7 \pm 2.1$ |
| F67A | Not detectable | Not detectable |
| D71N | $7.6 \pm 1.6$ | $18.4 \pm 8.0$ |
| D85N | $6.1 \pm 1.1$ | $32.9 \pm 26.8$ |
| H87A | $1.7 \pm 0.7$ | $21.9 \pm 6.4$ |
| H97A | $6.9 \pm 1.4$ | $5.4 \pm 0.7$ |
| F99A | $21.8 \pm 7.2$ | Not detectable |
| K101R | Not detectable | Not detectable |
| F116A | $7.0 \pm 0.7$ | $14.1 \pm 6.3$ |
| K193A | $2.0 \pm 0.8$ | $10.9 \pm 2.5$ |
| F67A/F99A | Not detectable | Not detectable |
| H87A/H97A | $1.8 \pm 1.7$ | $15.7 \pm 10.5$ |

The VqmA HTH region is enriched in positively charged residues. Within this region, we predicted that the residues K185, K193, and R195 are important for DNA binding due to their solvent accessibility and their conservation among VqmA orthologs (Fig. 4D). Indeed, as judged by EMSA analysis, changing any of these residues to alanine eliminated VqmA DNA binding (Fig. S4). This analysis is consistent with the structure of VqmA bound to DNA, which shows that residue K185 makes direct contact with the DNA backbone (17). We compared our structure of DPO-VqmA with a recently reported structure of DPO-VqmA-DNA (17). Alignment based on the LBD PAS domains reveals a conformational change of approximately 30 Å and a rotation of nearly 120° for the DBD (Fig. 4E). Thus, the DBD of the DNA-free VqmA protein adopts an open conformation, apparently conducive to DNA binding, while DNA-bound VqmA exhibits a closed conformation.

To probe the VqmA ligand binding requirements, we tested a panel of 15 synthesized or commercially available pyrazine, pyrimidine, and pyridine DPO analogs (Fig. 5A). We used the *V. cholerae* Δ*tdh* P*vqmR-lux* reporter strain and companion ITC analysis to, respectively, assess activation of and binding to VqmA (Fig. 5B and Table S3, respectively). Surprisingly, given the promiscuity VqmA displays with respect to DPO and Ala-AA binding, VqmA is highly selective in terms of cyclic compounds: VqmA responded most avidly to DPO, and the most potent analogs, capable of inducing maximal *lux* expression, had EC_50_ values 6-8-fold higher than that of DPO (Fig. 5C). Our results indicate that a 2’ hydroxyl or 2’ carbonyl group is necessary for binding to VqmA (Table S3) and promoting its transcriptional activity (Fig. 5C). Indeed, compound 10, which differs only from DPO by the lack of the 2’ hydroxyl group, was inactive and showed no binding to VqmA (Fig. 5B and Table S3).

**Figure 5.**
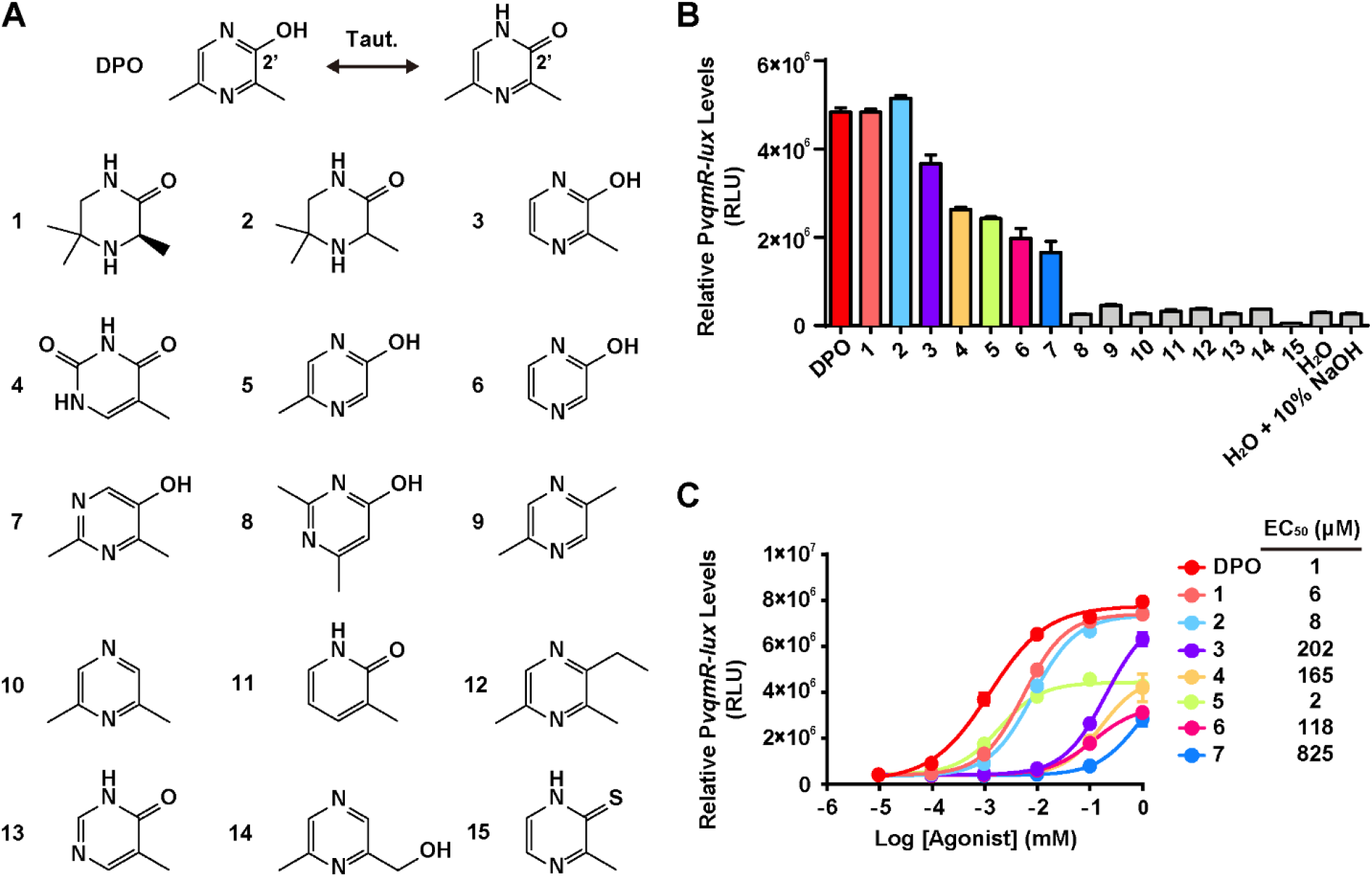
Structure-activity relationship study of DPO. (A) Structure of DPO and analogs studied here. (B) Responses of the P*vqmR-lux* reporter strain to 1 mM DPO and analogs. Compounds exhibiting >25% of the maximal activity (colored bars) and < 25% maximal activity (gray bars). (C) Dose response analyses of DPO and active analogs from (B) using the P*vqmR-lux* reporter strain. Dose responses are depicted as curve fits with the raw data plotted as individual points. Data points in (B) and (C) are represented as means ± SD with n = 3 biological replicates.

Our crystal structure shows that DPO binding is primarily mediated by π-π stacking forces contributed by residues F67 and F99 coupled with hydrogen bonding between the DPO 2’ carbonyl group and residue K101 (Fig. 4B). We note that DPO exists as the enol form in solution (8). Here, we built the keto tautomer of DPO in our crystal structure because it best fits the local density. Likely, the difference between the restricted condition in the crystal and the unrestricted environment in solution drives the preference for a particular tautomer. To verify our structure and to explore how VqmA could bind to both DPO and to Ala-AA, we conducted site-directed mutagenesis and tested binding and activity of DPO and Ala-AA, using an *in vitro* ITC assay and our *in vivo* cell-based reporter assay, respectively. We determined that, for WT Apo-VqmA, the K_d_ for DPO is ~2.3 μM, whereas the K_d_ for Ala-AA is ~14 μM (Fig. 6A). As a negative control, we performed ITC on a linear Ala-AA derivative called (2*S*)-2-amino-N-[(2*R*)-2-hydroxypropyl]propanamide (AHP) (Fig. S5A) that did not activate VqmA in our P*vqmR* reporter assay (Fig. S5B), and no binding was detected by ITC (Fig. S5C). Next, we focused on mutations in the DPO binding site: K23R, Y36F, F67A, D71N, D85N, H87A, H97A, F99A, K101R, F116A, F67A/F99A, and H87A/H97A (Fig. 4B). ITC results reveal that most of these VqmA variants exhibited approximately WT binding affinity for both DPO and Ala-AA (Table 1). Exceptions were VqmA F67A, VqmA K101R, and VqmA F67A/F99A, which showed no binding to either molecule indicating that these residues are crucial for binding to both DPO and Ala-AA (Table 1). Notably, VqmA F99A did not bind Ala-AA but retained residual binding to DPO (~10-fold higher K_d_ relative to WT), suggesting that the contribution of F99 in the π-π interaction with DPO is less crucial than that of F67. The finding that VqmA K101R was unable to bind DPO coupled with the structure-activity relationship analysis in Fig. 5, suggests that proper hydrogen bonding between K101 and the DPO 2’ hydroxyl/carbonyl group is critical for ligand binding and activity. We suspect that introduction of the bulkier Arg at position 101 prohibits formation of the required hydrogen bond at the 2’ position of DPO. A hydroxypyrimidine (compound 7, Fig. 5A) reduces the basicity of the oxygen on the 2’ moiety of the ligand, which would weaken its interaction with K101 relative to DPO. Indeed, compound 7 had an >800-fold increase in EC_50_ and exhibited no detectable binding to VqmA (see Fig. 5C and Table S3). Finally, the VqmA Y36F and H97A alterations made the K_d_ values for DPO and Ala-AA more similar, indicating that these residues are involved in dictating the natural preference VqmA shows for DPO over Ala-AA (Table 1).

**Figure 6.**
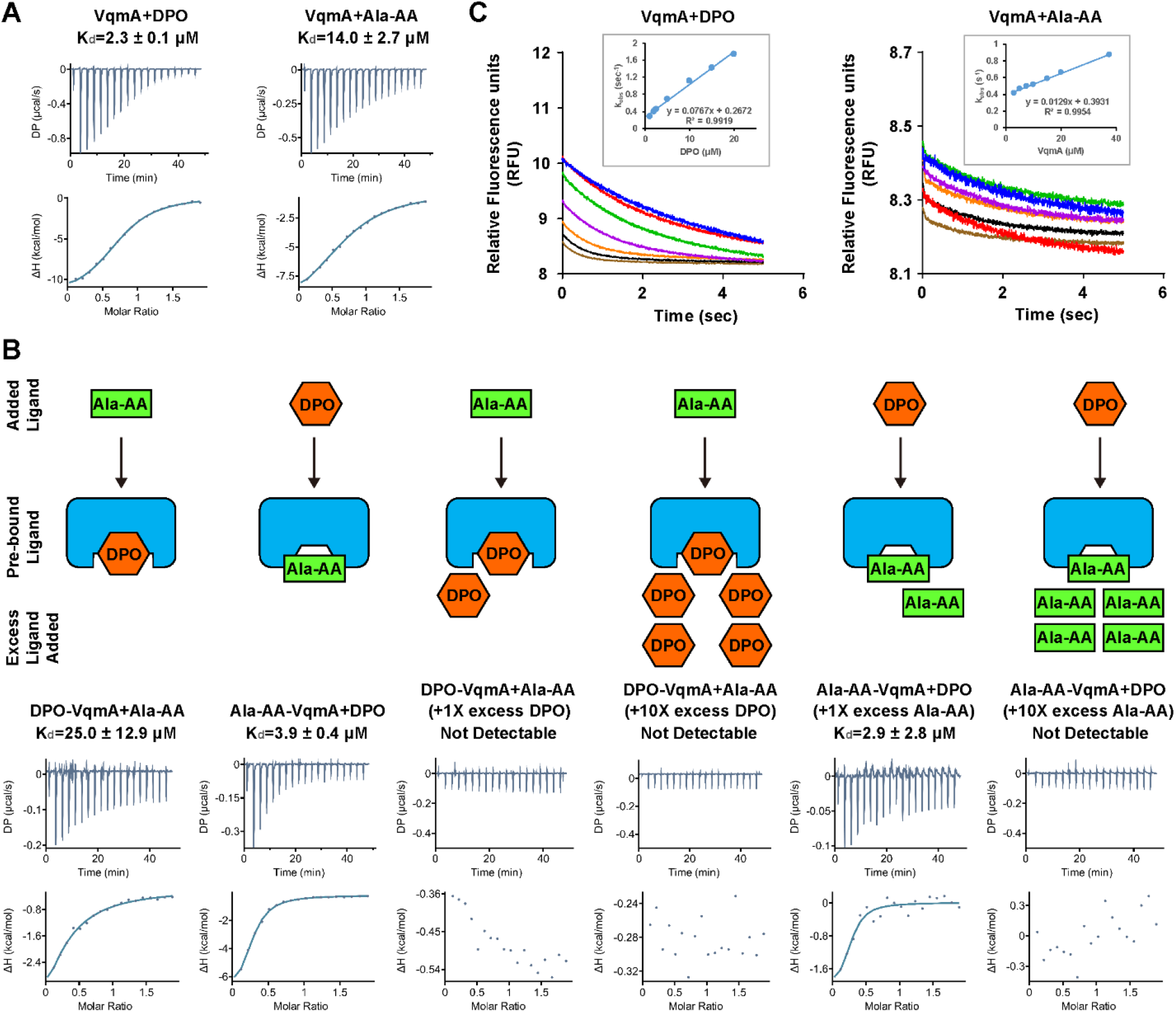
DPO and Ala-AA share the same binding site with DPO being the preferred ligand. (A) ITC measurement of the binding affinity of Apo-VqmA for DPO and Ala-AA. (B) ITC-based competition assay. Left to right: titration of Ala-AA into VqmA pre-bound with DPO, titration of DPO into VqmA pre-bound with Ala-AA, titration of Ala-AA into VqmA pre-bound to DPO with 1 -fold and 10-fold excess DPO present in solution, titration of DPO into VqmA pre-bound with Ala-AA with 1-fold and 10-fold excess Ala-AA present in solution. A cartoon schematic of each assay setup is shown above the corresponding ITC results. (C) Rates of DPO and Ala-AA binding to Apo-VqmA derived from stopped-flow binding assays. [Analyte] from low to high is blue, red, green, purple, orange, black, brown. See Methods for curve fitting in the insets.

The results from the reporter assay mimic those from the ITC assay (Fig. S6A). Fig. S6B shows that all of the VqmA variant proteins were produced at WT levels. Thus, changes in ligand-driven reporter activity stem from alterations in ligand binding affinity, not changes in protein stability. Our structure-guided mutagenesis and biochemical analyses suggest that DPO and Ala-AA occupy the same binding site on VqmA. To date, we have been unable to obtain crystal structures of Apo-VqmA or VqmA bound to Ala-AA. Apo-VqmA crystals diffract poorly and Ala-AA, while stable in solution as the HCl salt, tends to cyclize under our crystallization conditions independent of VqmA.

### Ala-AA and DPO share the same binding site on VqmA with DPO being the preferred ligand

Based on the surprising result that both Ala-AA and DPO bind to and activate VqmA, we sought to further characterize their interactions with VqmA to substantiate the same-binding-site hypothesis raised from our site-directed mutagenesis experiments. To garner additional evidence that DPO and Ala-AA bind the same site on VqmA, we performed an ITC-based competition assay. We titrated either DPO or Ala-AA into VqmA that had been pre-bound to the other molecule and assessed binding. We found that binding of the exogenously added molecule occurred no matter which ligand was pre-bound (Fig. 6B, left two panels), suggesting that the ~6-fold difference in K_d_ between the two ligands allows each molecule to displace the other. In a conceptually similar experiment, we performed the same titration in the presence of excess of the pre-bound ligand. Our rationale is that, if Ala-AA and DPO do share the same binding site in VqmA, the presence of an excess of the ligand that had been pre-bound should inhibit binding of the subsequently-added ligand. In this setup, VqmA showed a clear preference for binding to DPO. Specifically, when either an equal concentration or a 10-fold excess of DPO was provided to DPO-bound VqmA, Ala-AA was prevented from replacing pre-bound DPO (Fig. 6B, middle two panels). In the opposite case, only when there was at least 10-fold excess Ala-AA could Ala-AA prevent DPO from replacing pre-bound Ala-AA on VqmA (Fig. 6B, right two panels). These data support the hypothesis that DPO and Ala-AA share the same binding site on VqmA and further corroborate that, between the two ligands, DPO is preferred.

The preference VqmA shows for DPO over Ala-AA in the competitive binding experiments is curious given that the K_d_ values of the two ligands for the Apo-protein are only ~6-fold different. To better understand VqmA ligand binding preferences, we investigated the kinetics of DPO and Ala-AA binding using stopped-flow fluorimetry. When excited by 321 nm light, DPO emission occurs at 392 nm and the fluorescence is quenched upon binding to VqmA (Fig. S7A). Ala-AA is not fluorescent, but quenching of natural VqmA fluorescence (λ_ex_=280 nm/λ_em_=340 nm) can be monitored to track Ala-AA binding (Fig. S7B). We found that the k_on_ for DPO and Ala-AA are 7.7×10^4^ M^-1^·S^-1^ and 1.3×10^4^ M^-1^·S^-1^, respectively, while the calculated koff values are 0.27 S^-1^ and 0.39 S^-1^, respectively (Fig. 6C). Thus, the association rate of DPO is ~6-fold faster than that of Ala-AA while the dissociation rate of DPO is modestly slower, ~0.7-fold of that of Ala-AA. The calculated K_d_ (k_off_/k_on_) values from the stopped-flow experiments are 3.5 μM (DPO) and 30.5 μM (Ala-AA). This result is consistent with our ITC data because the differences are within an order of magnitude, which is well-accepted for these two methods. Moreover, the relative differences between the affinities of the two ligands for VqmA is consistent between the two methods. Taken together, we conclude that, once DPO is bound to VqmA, the complex does not easily dissociate, which prohibits subsequent binding of Ala-AA to the same binding site. In the reverse case, DPO can readily replace Ala-AA.

## DISCUSSION

In this study, we characterize ligand-controlled activation of VqmA, a cytoplasmic QS receptor and transcription factor. We show that VqmA is soluble, properly folded, and activates basal-level transcription of its target, *vqmR*, in the absence of its cognate AI, DPO. We show that the transcriptional activity of VqmA is increased in response to increasing concentrations of DPO, allowing VqmA to drive the *V. cholerae* QS transition to HCD upon DPO accumulation. A previous study reported that exogenous addition of DPO to purified Apo-VqmA does not increase the binding affinity of VqmA for P*vqmR* (17). We find that this is not the case as the DPO-bound VqmA protein has ~2-8-fold higher affinity for DNA than does Apo-VqmA (Fig. 1C, D). One possible explanation for this discrepancy is that the authors of the previous study purified VqmA from WT *E. coli*, which is *tdh*^+^. As mentioned, WT *E. coli* makes DPO and VqmA produced in *E. coli* binds to it (8). Thus, the authors’ putative Apo-VqmA could in fact have been Holo-VqmA. Consequently, their VqmA was likely saturated with DPO, and addition of more DPO would not enhance the VqmA DNA binding properties.

We contrast the behavior of the VqmA receptor to that of a canonical LuxR-type QS AI receptor-transcription factor that is activated only when the ligand is present (Fig. 2). While we cannot exclude issues arising from using different reporter systems, the dramatic difference in fold-change we observed between the VqmA and LasR receptors in the presence versus absence of their cognate AIs (3-fold for DPO/VqmA and 143-fold for 3OC_12_-HSL /LasR) leads us to speculate on the biological consequences of the differences between these systems. In particular, the ability of VqmA to function in both a ligand-independent and ligand-dependent manner enables the VqmA pathway to be active across all cell densities. In contrast, because LasR requires its AI for folding and activity, induction of the LasR pathway likely occurs exclusively under conditions when AI is present. We suggest that VqmA proteins are poised to rapidly respond to DPO when it appears. By contrast, LuxR-type receptors require accumulation of cognate AIs for proper folding and activation, therefore delaying QS activation to later in growth. Lastly, we note that while *V. cholerae vqmA* is expressed constitutively, *vqmA_Phage_* is not (10). While we do not yet know the natural inducer of *vqmA_Phage_*, we suspect that its production is regulated. We propose that the DPO-independent activity of VqmAPhage could allow the phage to overcome the requirement for the host to produce the AI and enable lysis provided that sufficient VqmAPhage is present.

The VqmA crystal structures reveal that the DPO binding pocket is located in the N-terminal PAS domain. Previous reports compared the VqmA fold to that of LasR, TraR, CviR, and other LuxR-type proteins (17,18), the latter of which, as mentioned, function via simultaneous binding and folding around AHL AIs (13,15,16). While this comparison is certainly informative, it is also worth noting that LuxR-type proteins do not contain PAS domains. PAS domains contain characteristic five stranded antiparallel β-sheet cores, while the LBDs of LuxR-type proteins contain four stranded antiparallel β-sheet cores. The divergence in folds is exemplified by the relatively low Z-scores of LuxR-type proteins compared to the VqmA structure (Table S2). We imagine that the LBDs of LuxR-type proteins constrain their selectivity to AHL molecules. By contrast, PAS domains are one of the most highly represented domains among signaling proteins and are capable of recognizing a wide range of ligands and sensory inputs, such as redox, light, gases, and metabolites (22–24). Within prokaryote taxa, however, it is unusual for a single PAS domain to bind multiple structurally diverse ligands (23). We are curious to discover what properties conferred by the VqmA PAS domain might enable the receptor to bind both the cyclic DPO and the linear Ala-AA molecules. Finally, despite the prevalence of PAS domains, VqmA represents a unique structure for QS receptors.

Our site-directed mutagenesis and ITC results indicate that DPO and Ala-AA use the same site in VqmA because the identical residues are required for binding and activation by both ligands. Confirmation comes from the competition and kinetic assays. We do note that the enthalpy changes measured in the ITC competition assay (Fig. 6B) are not the same as those that we measured for binding of DPO and Ala-AA to Apo-VqmA when there is no competition (Fig. 6A). While not investigated here, one possibility is that some Apo-VqmA remains present when we pre-bind VqmA to a ligand due to ligand association and disassociation. During the competition ITC, a portion of the ΔH value could stem from ligand binding to the small fraction of Apo-VqmA that is present. A second possibility is that ligand replacement in the binding pocket during the competition involves more than simply ligand exchange, for example protein breathing.

As noted, Ala-AA was a proposed intermediate in DPO production based on the knowledge that threonine and alanine are both substrates in DPO biosynthesis, and the proven structure of DPO (8). However, our present work indicates that Ala-AA is likely not the biological DPO precursor. Nonetheless, it is clear that Ala-AA is able to activate VqmA similar to DPO. It is not simply a matter of the VqmA receptor displaying ligand promiscuity, as we find VqmA is selective for DPO relative to other pyrazine analogs (Fig. 5). This result suggests that the activity conferred by Ala-AA is not coincidental, but likely the actual precursor to DPO is a molecule similar to Ala-AA. Identifying the biological precursor and exploring its role in VqmA activation are ongoing.

In addition to VqmA, other examples of proteins capable of binding multiple ligands exist, particularly those involved in metabolism. For example, IclR of *E. coli* binds pyruvate or glyoxylate, leading to activation or repression, respectively, of the operon regulating glyoxylate metabolism (25,26). This mechanism could enable a single receptor to measure glycolytic flux and efficiently adjust gene expression accordingly. Second, BenM of *Acinetobacter* binds benzoate and muconate, where muconate is a product of benzoate metabolism (27–29). Simultaneous binding of both molecules leads to transcriptional activation of the *benABCDE* operon, which further accelerates catabolism of benzoate to muconate (27–29). This arrangement enables *Acinetobacter* to tightly control the expression of genes required for the degradation of benzoate, only activating them when their functions are required. Distinct from both of these cases, VqmA binds to either DPO or Ala-AA, but not both, to induce maximal transcriptional activity. This result is curious to us given that QS receptors typically maintain signaling fidelity by distinguishing between different ligands (30–33). This situation appears analogous to that of the *V. cholerae* AI called CAI-1 ((*S*)-3-hydroxytridecan-4-one) and its biosynthetic precursor, Ea-CAI-1 (3-aminotridec-2-en-4-one). The study of Wei *et al*. showed that Ea-CAI-1 is a potent activator of CqsS, but this precursor is likely more sensitive to hydrolysis than CAI-1, and therefore is too unstable to be a satisfactory QS signal (34). Given that pyrazine molecules are known to be stable compounds (35) and Ala-AA tends to cyclize, we hypothesize that while DPO is not an absolute requirement for VqmA-mediated QS, DPO could be the more optimal AI relative to Ala-AA because of its stability and potency.

Finally, we do note that all three *V. cholerae* QS systems converge to control an overlapping group of targets, including the *aphA* gene encoding the QS master regulator AphA (6,9) that is expressed at LCD when AI concentrations are below their detection thresholds. The HCD responses to CAI-1, AI-2 (4,5-dihydroxy-2,3-pentanedione) (36,37), and the DPO (8) AIs drive repression of *aphA*. VqmA, because it does not strictly require an AI for activity, activates some *vqmR* expression at LCD, leading to some repression of *aphA*. Thus, VqmA-controlled QS could provide a mechanism to attenuate AphA production, and thus alter expression of its downstream targets, prior to when sufficient levels of CAI-1, AI-2, and DPO have accumulated for the transition to the HCD QS state. Subsequently, DPO-VqmA-mediated QS could reinforce repression of *aphA* once *V. cholerae* has committed to the HCD QS program. Thus, the VqmA QS circuit could be important, at all cell densities, for fine-tuning the expression of QS target genes, and in turn, collective behaviors.

## EXPERIMENTAL PROCEDURES

### Bacterial strains, plasmids, primers, and reagents

Strains, plasmids, and primers used in this study are listed in Tables S4-6, respectively. Unless otherwise indicated, *V. cholerae* and *E. coli* were grown aerobically in LB at 37°C. M9 medium was supplemented with 0.5% glucose and the indicated amino acids, each at 0.4 mM. Antibiotics were used at the following concentrations: 50 U mL^−1^ polymyxin B, 200 μg mL^−1^ ampicillin, and 100 μg mL^-1^ kanamycin. Primers were obtained from Integrated DNA Technologies. Site directed mutagenesis of *vqmA* was performed using PCR with the indicated primers and the cloning strategy described in Table S6. The same primer pairs were used for generating mutants in both the *E. coli* expression vector (JSS-1057) and the *V. cholerae* arabinose-inducible vector (KPS-0500). Intra-molecular reclosure was performed as described (10) using Q5 High-Fidelity DNA Polymerase (NEB). QuickChange (Agilent) site directed mutagenesis was performed as described (30) using PfuUltra Polymerase (Agilent). The parent pET15b-*vqmA* vector (JSS-1057) was constructed by PCR of pET15b-*His-vqmA* (KPS-0929) with primers JSO-654 x JSO-655, followed by intramolecular-reclosure to remove the His-tag. The P*vqmR-lux::lacZ* suicide vector (JSS-0807) was constructed with primers JSO-383 x JSO-384 on BB-Ec0361 (pCN004) to generate a one-sided PacI-digestable pKAS backbone with flanking *lacZ* sites. The P*vqmR-lux* insert was generated using primers JSO-358 x JSO-397 on JSS-743 (P*vqmR-lux* in a pEVS143-derived vector). The insert and vector were digested with PacI (NEB), ligated using T4 DNA ligase (NEB), and transformed into *E. coli* S17-1 λ*pir*.

### Molecule syntheses

Unless otherwise indicated, all temperatures are expressed in °C (degrees Centigrade). All reactions were conducted at room temperature unless otherwise noted. ^1^H-NMR spectra were recorded on a Varian VXR-400, or a Varian Unity-400 at 400MHz field strength. Chemical shifts are expressed in parts per million (ppm, δ units). Coupling constants (*J*) are in units of hertz (Hz). Splitting patterns describe apparent multiplicities and are designated as s (singlet), d (doublet), t (triplet), q (quartet), m (multiplet), quin (quintet) or br (broad). The mass spec was run on a Sciex API 100 using electrospray ionization (ESI). The LC-MS was run using a C-18 reverse phase column (2.1 ID, 3.5 micron, 50 mm). The column conditions were 98% water with 0.05%TFA and 2% MeOH to 100% MeOH over 5.5 minutes. Analytical thin layer chromatography was used to verify the purity as well as to follow the progress of reaction(s). Unless otherwise indicated, all final products were at least 95% pure as judged by HPLC / MS.

### (*S*)-1-oxo-1-((2-oxopropyl)amino)propan-2-aminium chloride (Ala-AA)

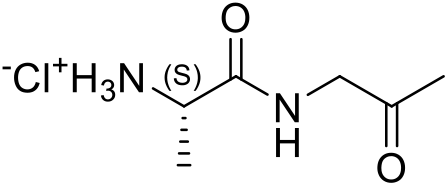

Step 1: *tert*-butyl ((*S*)-1-(((*R*)-2-hydroxypropyl)amino)-1-oxopropan-2-yl)carbamate

A solution of (*tert*-butoxycarbonyl)-*L*-alanine (20.0 g, 105 mmol, 1.0 eq) in DCM (50 mL) was stirred and cooled to 0°C, and then 4-methylmorpholine (12.8 g, 126 mmol, 13.9 mL, 1.20 eq) was added. The resulting solution was stirred 5 min, then isobutyl chloroformate (15.8 g, 116 mmol, 15.2 mL, 1.10 eq) was added dropwise over 15 min. The resulting reaction mixture was stirred at 0°C for 1 hr, then (*R*)-1-aminopropan-2-ol (8.34 g, 110 mmol, 8.74 mL, 1.05 eq) was added drop-wise over 10 min. The resulting reaction mixture was allowed to warm to RT and stirred for 12 hr. The reaction mixture was concentrated under reduced pressure. The residue was purified by silica gel column chromatography (DCM:MeOH = 100:0 to 98:2) to afford *tert*-butyl ((*S*)-1-(((*R*)-2-hydroxypropyl)amino)-1-oxopropan-2-yl)carbamate (13.2 g, 48% yield, 95% purity) as a yellow oil. ^1^H NMR (400 MHz, CDCl_3_) δ = 6.83 (s, br, 1H), 5.24 (s, br, 1H), 4.15 (m, 1H), 3.91 (m, 1H), 3.38 (m, 1H), 3.12 (m, 2H), 1.46 (s, 9H), 1.36 (d, *J* = 7.2, 3H), 1.17 (d, *J* = 6.4, 3H).

Step 2: *tert*-butyl (*S*)-(1-oxo-1-((2-oxopropyl)amino)propan-2-yl)carbamate

*Tert*-butyl ((*S*)-1-(((*R*)-2-hydroxypropyl)amino)-1-oxopropan-2-yl)carbamate (8.00 g, 32.4 mmol, 1.00 eq) was dissolved in DCM (65 mL) and cooled to 0°C. Next, Dess-Martin periodinane (17.9 g, 42.2 mmol, 13.1 mL, 1.30 eq) was added, and the resulting reaction mixture was warmed to RT and stirred for 12 hr. The reaction mixture was quenched by addition of saturated aqueous Na_2_S_2_O_3_ (100 mL) and saturated aqueous NaHCO_3_ solution (100 mL), then stirred for 30 min and filtered through a pad of Celite. The filtrate was extracted with DCM (3 x 100 mL). The combined organic layers were washed with brine (100 mL), dried over anhydrous Na_2_SO_4_, filtered, and concentrated under reduced pressure to give a residue. The residue was purified by silica gel column chromatography (DCM:EtOAc = 100:0 to 20:1) to afford *tert*-butyl (*S*)-(1-oxo-1-((2-oxopropyl)amino)propan-2-yl)carbamate (3.70 g, 42% yield, 90% purity) as a yellow solid. ^1^H NMR (400 MHz, CDCl_3_) δ = 6.91 (s, br, 1H), 5.12, (s, br, 1H), 4.25 (m, 1H), 4.13 (d, *J* = 4.0, 2H), 2.19 (s, 3H), 1.44 (s, 9H), 1.36 (d, *J* = 7.2, 3H).

Step 3: (*S*)-1-oxo-1-((2-oxopropyl)amino)propan-2-aminium chloride

*Tert*-butyl (*S*)-(1-oxo-1-((2-oxopropyl)amino)propan-2-yl)carbamate (3.70 g, 15.2 mmol, 1.00 eq) was dissolved in HCl/dioxane (4.00 M, 18.9 mL, 5.00 eq) and the resulting reaction mixture was stirred at RT for 2 hr, during which time a precipitate appeared. The reaction mixture was filtered to remove liquids, and the filter cake was washed with MTBE (2 x 20 mL) and dried under vacuum. The crude solid was triturated with EtOAc (50 mL) at RT for 2 hr followed by filtration and trituration with MBTE (20 mL) at RT for 2 hr to afford (*S*)-1-oxo-1-((2-oxopropyl)amino)propan-2-aminium chloride (0.50 g, 17 % yield, 95 % purity) as a yellow solid. ^1^H NMR (400 MHz, D2O) δ = 4.75 (m, 1H), 4.12 (m, 1H), 2.12 (s, 3H), 1.41 (d, *J* = 6.4, 3H).

### (*S*)-1-oxo-1-((2-oxopropyl)amino)propan-3,3,3-d_3_-2-aminium chloride (d_3_-Ala-AA)

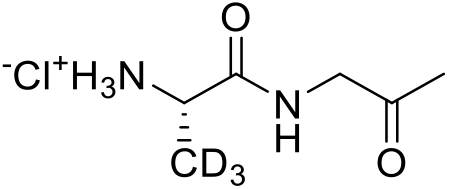

Step 1: *tert*-butyl ((*S*)-1-(((*R*)-2-hydroxypropyl)amino)-1-oxopropan-2-yl-3,3,3-d_3_)carbamate

A solution of (*tert*-butoxycarbonyl)-*L*-D_3_-alanine (2.0 g, 10.6 mmol, 1.0 eq) in DCM (20 mL) was stirred and cooled to 0°C, and then 4-methylmorpholine (1.28 g, 12.7 mmol, 1.39 mL, 1.20 eq) was added. The resulting solution was stirred 5 min, then isobutyl chloroformate (1.58 g, 11.6 mmol, 1.52 mL, 1.10 eq) was added dropwise over 15 min. The resulting reaction mixture was stirred at 0°C for 1 hr, then (*R*)-1-aminopropan-2-ol (834 mg, 11.0 mmol, 870 μL, 1.05 eq) was added drop-wise over 2 min. The resulting reaction mixture was allowed to warm to RT and stirred for 12 hr. The reaction mixture was concentrated under reduced pressure. The residue was purified by silica gel column chromatography (DCM:MeOH = 98:2) to afford *tert*-butyl ((*S*)-1-(((*R*)-2-hydroxypropyl)amino)-1-oxopropan-2-yl-3,3,3-d_3_)carbamate (2.0 g, 68% yield, 90% purity) as a colorless oil. ^1^H NMR (400 MHz, CDCl_3_) δ = 6.66 (s, br, 1H), 5.06 (s, br, 1H), 4.13 (m, 1H), 3.94 (m, 1H), 3.73 (m, 1H), 3.49, (m, 1H), 3.12 (m, 1H), 1.45 (s, 9H), 1.18 (d, *J* = 6.4, 3H).

Step 2: *tert-*butyl (*S*)-(1-oxo-1-((2-oxopropyl)amino)propan-2-yl-3,3,3-d_3_)carbamate

*Tert*-butyl ((*S*)-1-(((*R*)-2-hydroxypropyl)amino)-1-oxopropan-2-yl-3,3,3-d_3_)carbamate (2.00 g, 7.31 mmol, 1.00 eq) was dissolved in DCM (25 mL) and cooled to 0°C. Next, Dess-Martin periodinane (4.0 g, 9.5 mmol, 2.94 mL, 1.30 eq) was added, and the resulting reaction mixture was warmed to RT and stirred for 12 hr. The reaction mixture was quenched by addition of saturated aqueous Na_2_S_2_O_3_ (20 mL) and saturated aqueous NaHCO_3_ solution (20 mL), then stirred for 30 min and filtered through a pad of Celite. The filtrate was extracted with DCM (3 x 20 mL). The combined organic layers were washed with brine (20 mL), dried over anhydrous Na_2_SO_4_, filtered, and concentrated under reduced pressure to give a residue. The residue was purified by silica gel column chromatography (DCM:EtOAc = 98:2) to afford *tert-*butyl (*S*)-(1-oxo-1-((2-oxopropyl)amino)propan-2-yl-3,3,3-d_3_)carbamate (1.60 g, 81% yield, 90% purity) as a yellow solid. ^1^H NMR (400 MHz, CDCl_3_) δ = 6.82 (s, br, 1H), 4.99, (s, br, 1H), 4.21 (m, 1H), 4.15 (d, *J* = 4.0, 2H), 2.21 (s, 3H), 1.46 (s, 9H).

Step 3: (*S*)-1-oxo-1-((2-oxopropyl)amino)propan-3,3,3-d_3_-2-aminium chloride

*Tert*-butyl (*S*)-(1-oxo-1-((2-oxopropyl)amino)propan-2-yl-3,3,3-d_3_)carbamate (1.60 g, 5.89 mmol, 1.00 eq) was dissolved in HCl/dioxane (4.00 M, 7.37 mL, 5.00 eq) and the resulting reaction mixture was stirred at RT for 2 hr, during which time a yellow precipitate appeared. The reaction mixture was filtered to remove liquids, and the filter cake was washed with MTBE (2 x 20 mL) and dried under vacuum. The crude solid was triturated with EtOAc (50 mL) at RT for 2 hr followed by filtration and trituration with MBTE (20 mL) at RT for 2 hr to afford (*S*)-1-oxo-1-((2-oxopropyl)amino)propan-3,3,3-d_3_-2-aminium chloride (0.67 g, 60 % yield, 95 % purity) as a yellow solid. ^1^H NMR (400 MHz, D2O) δ = 4.75 (s, 2H), 4.15 (s, 1H), 2.16 (s, 3H).

### (*S*)-2-amino-N-((*R*)-2-hydroxypropyl)propanamide (AHP)

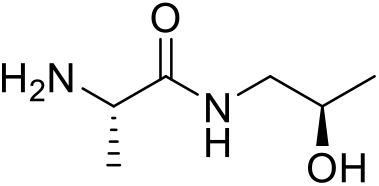

Step 1: Benzyl ((*S*)-1-(((*R*)-2-hydroxypropyl)amino)-1-oxopropan-2-yl)carbamate

(2*S*)-2-(benzyloxycarbonylamino)propanoic acid (5 g, 22.40 mmol, 1.0 eq) was dissolved in THF (100 mL) and the resulting solution was cooled to 0 °C under an N_2_ atmosphere. Next, 4-methylmorpholine (4.53 g, 44.80 mmol, 4.93 mL, 2.0 eq) was added, followed by isobutyl chloroformate (3.06 g, 22.40 mmol, 2.94 mL, 1.0 eq). The resulting reaction mixture was stirred at 0 °C for 1 hr. Then (2R)-1-aminopropan-2-ol (2.52 g, 33.60 mmol, 2.65 mL, 1.5 eq) was added to the reaction mixture, and the resulting mixture was allowed to warm to RT at stirred 11 hr. The reaction was concentrated under reduced pressure to remove THF. The reaction mixture was diluted with H_2_O (50 mL), then extracted with EtOAc (2 x 30 mL). The combined organic layers were dried over Na_2_SO_4_, filtered, and concentrated under reduced pressure to give a residue. The residue was purified by silica column chromatography (PE:EtOAc = 1:1 to 0:100) to afford benzyl ((*S*)-1-(((*R*)-2-hydroxypropyl)amino)-1-oxopropan-2-yl)carbamate (3.0 g, 48% yield) as a white solid. ^1^H NMR (400 MHz, CDCl_3_) δ = 7.35 (m, 5H), 6.79 (s, br, 1H), 5.62 (d, br, *J* = 6.0, 1H), 5.10 (m, 2H), 4.24 (m, 1H), 3.88 (s, br, 1H), 3.36 (m, 1H), 3.12 (m, 2H), 1.37 (d, *J* = 7.2, 3H), 1.15 (d, *J* = 6.0, 3H); LC-MS calculated for C_14_H_20_N_2_O_4_: *m/z* = 280; found: *m/z* = 281 (M+H).

Step 2: (*S*)-2-amino-N-((*R*)-2-hydroxypropyl)propanamide

To a solution of benzyl ((*S*)-1-(((*R*)-2-hydroxypropyl)amino)-1-oxopropan-2-yl)carbamate (1 g, 3.57 mmol, 1.0 eq) in MeOH (10 mL) was added Pd(OH)_2_/C (1 g, 20% by weight) under N_2_. The suspension was degassed under vacuum and purged with H_2_ several times. The mixture was stirred under H_2_ (15 psi) at RT for 2 hr. The reaction mixture was flushed 3x with N_2_ to degas then filtered through a pad of Celite to remove the Pd, and the filter cake was washed with MeOH (3 x 10 mL). The filtrate was concentrated under reduced pressure to give a residue. The residue was purified by alumina column chromatography (EtOAc:MeOH = 100:0 to 10:1) to afford (*S*)-2-amino-N-((*R*)-2-hydroxypropyl)propanamide (500 mg, 95% yield) as a white solid. ^1^H NMR (400 MHz, MeOH-d_4_) δ = 3.83 (m, 1H), 3.42 (q, *J* = 6.8, 1H), 3.24 (dd, *J* = 4.8, 8.8, 1H), 3.12 (dd, *J* = 6.4, 8.8, 1H), 1.27 (d, *J* = 6.8, 3H), 1.15 (d, *J* = 6.4, 3H); LC-MS calculated for C_6_H_14_N_2_O_2_: *m/z* = 146; found: *m/z* = 147 (M+H).

### (*R*)-3,5,5-trimethylpiperazin-2-one (compound 1)

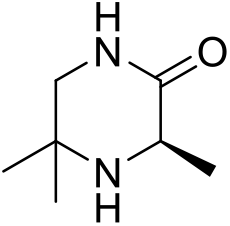

Step 1: Methyl (2-methyl-1-(tritylamino)propan-2-yl)-*L*-alaninate

2-methyl-N1-trityl-propane-1,2-diamine (2.2 g, 6.66 mmol, 1.0 eq) and Et_3_N (1.01 g, 9.99 mmol, 1.39 mL, 1.5 eq) were dissolved in DCM (50 mL) and cooled to 0 °C. Next, methyl (2*S*)-2-(trifluoromethyl-sulfonyloxy)propanoate (1.57 g, 6.66 mmol, 1.0 eq) was added, and the resulting reaction mixture was allowed to stir for 1 hr at 0 °C and then warmed to RT and stirred for 12 hr. The reaction mixture was diluted with NaHCO_3_ (20 mL) and extracted with EtOAc (3 x 20 mL). The combined organic layers were dried over Na_2_SO_4_, filtered, and concentrated under reduced pressure to give a residue. The residue was purified by silica gel column chromatography (PE:EtOAc = 100:1 to 50:1) to afford methyl (2-methyl-1-(tritylamino)propan-2-yl)-*L*-alaninate (1.4 g, 50% yield) as a colorless oil. ^1^H NMR (400 MHz, CDCl_3_) δ = 7.46 (d, *J* = 7.5, 6H), 7.23 (m, 6H), 7.13 (m, 3H), 3.55 (s, 3H), 2.90 (q, *J* = 7.1, 1H), 1.95 (m, 2H), 1.09 (s, 3H), 1.04 (d, *J* = 7.1, 3H), 0.96 (m, 3H).

Step 2: (*R*)-3,5,5-trimethylpiperazin-2-one

A solution of methyl (2-methyl-1-(tritylamino)propan-2-yl)-*L*-alaninate (1.2 g, 2.88 mmol, 1.0 eq) in DCM (7.5 mL) was cooled to 0 °C and then TFA (5 mL) was added dropwise. The resulting reaction mixture was warmed to RT and stirred for 2 hr. The mixture was concentrated under reduced pressure to afford a residue. The residue was diluted with NaOH (10%, 20 mL), then extracted with DCM (3 x 30 mL). The combined organic layers were dried over Na_2_SO_4_, filtered, and concentrated under reduced pressure to give a residue. The residue was purified by silica gel column chromatography (PE:EtOAc = 1:1 to 0:100) to afford a yellow solid. The solid was triturated for 10 min with n-pentane (10 mL), filtered, and dried under vacuum to afford (*R*)-3,5,5-trimethylpiperazin-2-one (180 mg, 44% yield) as a light yellow solid. ^1^H NMR (400 MHz, CDCl_3_) δ = 5.92 (s, br, 1H), 3.62 (q, *J* = 6.8, 1H), 3.29 (d, *J* = 11.6, 1H), 3.08 (dd, *J* = 4.5, 11.6, 1H), 1.39 (d, *J* = 6.8, 3H), 1.31 (s, 3H), 1.21 (s, 3H); LC-MS calculated for C_7_H_14_N_2_O: *m/z* = 142; found: *m/z* = 143 (M+H).

### 3,5,5-trimethylpiperazin-2-one (compound 2)

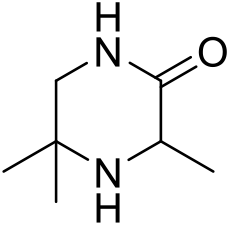

Step 1: Methyl 2-(trifluoromethylsulfonyloxy)propanoate

A solution of methyl 2-hydroxypropanoate (5.0 g, 48.03 mmol, 4.59 mL, 1.0 eq) and 2,6-dimethylpyridine (5.15 g, 48.03 mmol, 5.59 mL, 1.0 eq) in DCM (50 mL) was cooled to −10 °C, degassed and purged with N_2_ for 3 times, then trifluoromethylsulfonyl trifluoromethanesulfonate (16.26 g, 57.63 mmol, 9.51 mL, 1.2 eq) was added dropwise at −10 °C under N_2_. The resulting reaction mixture was stirred at −10 °C for 30 min under N_2_ atmosphere. The reaction mixture was quenched by addition of aqueous hydrochloric acid (0.5 M, 50 mL) and stirred at − 10°C for another 30 min, then the reaction mixture was transferred to a separatory funnel and the layers were separated. The organic layer was washed with brine (30 mL), dried over Na_2_SO_4_, filtered and concentrated under reduced pressure to give a residue. The residue was purified by silica gel column chromatography (PE:EtOAc = 100:0 to 10:1) to afford methyl 2-(trifluoromethylsulfonyloxy)propanoate (6.03 g, 53% yield) as a brown oil. ^1^H NMR (400 MHz, CDCl_3_) δ = 5.25 (q, *J* = 7.0, 1H), 3.87 (s, 3H), 1.73 (d, *J* = 7.0, 3H).

Step 2: Methyl (2-methyl-1-(tritylamino)propan-2-yl)-alaninate

2-methyl-N1-trityl-propane-1,2-diamine (5.0 g, 15.13 mmol, 1.0 eq) and Et_3_N (1.53 g, 15.13 mmol, 2.11 mL, 1.0 eq) were dissolved in DCM (50 mL) and cooled to 0 °C. Next, methyl 2-(trifluoromethyl-sulfonyloxy)propanoate (3.57 g, 15.13 mmol, 1.0 eq) was added, and the resulting reaction mixture was allowed to stir for 1 hr at 0 °C and then warmed to RT and stirred for 6 hr. The reaction mixture was diluted with NaHCO_3_ (50 mL) and extracted with EtOAc (3 x 30 mL). The combined organic layers were dried over Na_2_SO_4_, filtered, and concentrated under reduced pressure to give a residue. The residue was purified by silica gel column chromatography (PE:EtOAc = 100:1 to 10:1) to afford methyl (2-methyl-1-(tritylamino)propan-2-yl)-alaninate (4.3 g, 61% yield) as a white solid. ^1^H NMR (400 MHz, MeOH-d_4_) δ = 7.48 (m, 6H), 7.26 (m, 6H), 7.17 (m, 3H), 3.59 (s, 3H), 3.05 (q, *J* = 7.2, 1H), 2.04 (m, 1H), 1.96 (m, 1H), 1.11 (d, *J* = 7.2, 3H), 1.08 (s, 3H), 1.03 (s, 3H).

Step 3: 3,5,5-trimethylpiperazin-2-one

A solution of methyl (2-methyl-1-(tritylamino)propan-2-yl)-alaninate (3.0 g, 7.2 mmol, 1.0 eq) in DCM (15 mL) was cooled to 0 °C and then TFA (10 mL) was added dropwise. The resulting reaction mixture was warmed to RT and stirred for 2 hr. The mixture was concentrated under reduced pressure to afford a residue. The residue was diluted with NaOH (10%, 20 mL), then extracted with DCM (3 x 30 mL). The combined organic layers were dried over Na_2_SO_4_, filtered, and concentrated under reduced pressure to give a residue. The residue was purified by silica gel column chromatography (PE:EtOAc = 1:1 to 0:100) to afford a yellow solid. The solid was triturated for 10 min with n-pentane (10 mL), filtered, and dried under vacuum to afford 3,5,5-trimethylpiperazin-2-one (168mg, 16% yield) as a white solid. ^1^H NMR (400 MHz, CDCl_3_) δ = 6.11 (s, br, 1H), 3.61 (q, *J* = 6.8, 1H), 3.27 (d, *J* = 11.6, 1H), 3.08 (dd, *J* = 4.5, 11.6, 1H), 1.39 (d, *J* = 6.8, 3H), 1.31 (s, 3H), 1.19 (s, 3H); LC-MS calculated for C7H14N_2_O: *m/z* = 142; found: *m/z* = 143 (M+H).

### Bioluminescence assays

Assays were performed as described (8) with the following modifications: *V. cholerae* reporter strains were grown overnight and back-diluted to OD_600_ = 0.005 in M9 medium supplemented with threonine and leucine. 150 μL of culture was transferred to the wells of a 96-well plate, followed by addition of 50 μL of cell-free culture fluids. Synthetic molecule standards were prepared in M9 medium, and 50 μL of these preparations were added to the *V. cholerae* reporter strains in place of cell-free fluids. AI dose response assays were performed as described (10) with the following modifications: 198 μL aliquots of cell cultures were dispensed into a 96-well plate with 2 μL aliquots of serially-diluted synthetic compounds. Control wells received 2 μL of solvent (water or 10% NaOH).

### Electromobility Shift Assays (EMSAs)

EMSAs were performed as described (10). If not specified otherwise, the highest concentration of VqmA assessed was 130 nM. Two-fold serial-dilutions of protein were used to generate lower concentrations. Biotinylated P*vqmR* DNA (540 pM) was used in all EMSAs. The percent of P*vqmR* DNA bound was calculated using the gel analyzer tool in ImageJ.

### Production of cell-free translated protein

Cell-free translated His-VqmA was produced using the S30 T7 High-Yield Protein Expression System (Promega) according to the manufacturer’s protocol with the following amendments: 1.5 μg purified plasmid DNA, encoding His-VqmA under the T7 promoter, was used as the template; 40 U of Murine RNase inhibitor (NEB) was added; 5 reactions were pooled together; and the reactions were incubated (translated) for up to 1.5 h at 37°C. After translation, the His-VqmA was purified using MagneHis Ni-Particles (Promega).

### VqmA/P*vqmR*, LasR/P*lasB*, and VqmA_Phage_/P*qtip* reporter assays

For each system being tested, Δ*tdh E. coli* harboring one plasmid encoding the arabinose-inducible receptor-transcription factor (VqmA, LasR, or VqmA_Phage_) and a second plasmid containing the cognate target promoter (P*vqmR*, P*lasB*, or P*qtip*, respectively) fused to *lux* (10,38) was used. In parallel, a control strain for each system was used that carried the reporter plasmid alone (P*vqmR-lux*, P*lasB-lux*, or P*qtip-lux*). Each strain was grown overnight and back diluted 1:500 into fresh LB containing 0.075% arabinose and either synthetic AI (20 μM DPO or 1 μM 3OC_12_-HSL) or an equivalent volume of solvent (water for DPO and DMSO for 3OC_12_-HSL). 200 μL aliquots of each culture were dispensed into 96-well plates and the plates were incubated at 30°C with shaking. OD_600_ and bioluminescence were measured at regular intervals for 20 h. The bars presented in Figs. 2 and S1C represent the timepoints with the largest differences in relative light units (RLU) between the AI-receiving and non-receiving samples for each system. The same timepoints were used for the reporter-only strains, which received both arabinose and AI.

### DPO extraction from cell-free culture fluids

*V. cholerae* strains were grown overnight and back-diluted to OD_600_ = 0.005 in M9 medium supplemented with leucine. The cultures were grown to OD_600_ = ~ 1.0, followed by incubation with 100 μM synthetic DPO or 100 μM synthetic d_3_-Ala-AA for 15 min. DPO, d_3_-DPO and d_3_-Ala-AA were extracted from cell-free culture fluids by SPE (HyperSep Hypercarb SPE cartridges, Thermo). Chromatographic separation was achieved with increasing concentrations of acetonitrile (0%, 10%, 20%, 30%, 50%, and 100%). The relative quantities of DPO, d_3_-DPO, and d_3_-Ala-AA were determined by LC-MS (described below). Samples were back-diluted 1:10 in water prior to column injection. Standards containing synthetic DPO, Ala-AA, or d_3_-Ala-AA were prepared in 2% acetonitrile.

### Ala-AA conversion to DPO

10 μM synthetic Ala-AA or 10 μM synthetic DPO was incubated with 50 μM Apo-VqmA in EMSA buffer. Each single component was also diluted into EMSA buffer as a control. Following incubation for 15 min, excess and non-protein-bound small molecules were removed by size-exclusion chromatography (Zeba), and the resulting protein was heated to 70°C for 10 min. Denatured protein was removed by centrifugation (13,000 x g, 10 min) and supernatants were assessed for Ala-AA or DPO by LC-MS (described below).

### Liquid Chromatography-Mass Spectrometry (LC-MS) to detect DPO, Ala-AA, d_3_-DPO, and d_3_-Ala-AA

LC-MS was performed with modifications (10). Samples and standards were loaded onto a C_18_ column (1 mm by 100 mm; ACE 3 C18 AQ, Mac-Mod) using a Shimadzu HPLC system and PAL auto-sampler (20 μL per injection) at a flow rate of 70 μL min^-1^. Full scan MS data were acquired with an LTQ-Orbitrap XL mass spectrometer (Thermo) at a resolution of 60,000 in profile mode from the *m/z* range of 110 to 200. DPO, Ala-AA, d_3_-DPO, and d_3_-Ala-AA were detected by performing XIC at *m/z* values of 125.0714 (C_6_H_9_N_2_O), 145.0977 (C_6_H_13_N_2_O_2_), 128.0903 (C_6_H_6_d_3_N_2_O), and 148.1165 (C_6_H_10_d_3_N_2_O_2_), respectively, each with a mass accuracy of ±10 ppm. Caffeine (2 μM in 50% acetonitrile with 0.1%formic acid) was delivered as a lock mass using an HPLC pump (LC Packing) with a flow splitter and at an effective flow rate of 20 μL min^-1^ through a tee at the column outlet. Raw files were imported into Skyline v4.2.0 (39) and peak areas for each molecule were extracted using the small molecule workflow.

### Protein expression and purification

*V. cholerae vqmA* was cloned into pET15b and overexpressed in Δ*tdh* BL21 (DE3) *E. coli* cells using 1 mM IPTG at 37°C for 4 h. Cells were pelleted at 16,100 x g for 15 min and resuspended in lysis buffer (20 mM Tris-HCl pH 8, 150 mM NaCl, 1 mM DTT). Cells were lysed using sonication and subjected to centrifugation at 32,000 x g for 30 min. The supernatant was loaded onto a heparin column (GE Healthcare) and eluted by a linear gradient from buffer A (20 mM Tris-HCl pH 8, 1 mM DTT) to buffer B (20 mM Tris-HCl pH 8, 1 M NaCl, 1 mM DTT). Peak fractions were pooled, concentrated, and loaded onto a Superdex-200 size exclusion column (GE Healthcare) in gel filtration buffer (20 mM Tris-HCl pH 8, 150 mM NaCl, and 1 mM DTT). Proteins were concentrated to 30 mg/mL, flash frozen, and stored at −80°C.

### Crystallization and structure determination

The DPO-VqmA complex was formed by mixing 1 mM purified Apo-VqmA with 10-fold excess synthetic DPO prior to crystallization. Crystals were obtained by the hanging drop vapor diffusion method from drops containing a 1:1 mixture of protein:precipitant (0.1 M Tris-HCl pH 7.3, 16% PEG3350 and 0.2 M Sodium Malonate) at 22°C. The trays were seeded using cat whiskers (Spark Bassler). Crystals were cryoprotected in a solution containing the precipitant plus 10% (v/v) glycerol. Crystals belonged to the P 2_1_2_1_2_1_ space group with unit cell dimensions a = 50.4 Å, b = 84.6 Å, c = 111.6 Å. A dataset with heavy atom derivatives was obtained by soaking the crystals in cryoprotectant supplemented with 10 mM KAu(CN)_2_ (Hampton Research) for 10 min at room temperature (RT) as reported (40). Crystals were subsequently flash frozen and stored in liquid N_2_. Diffraction data were collected at the NSLS II (National Synchrotron Light Source) beamlines FMX and AMX. Data were processed using HKL-3000 and experimental phases were determined using single-wavelength anomalous dispersion (41). Experimental phases were calculated using SHELX (42). The data sets were subjected to density modification using PHENIX (43–45) and the VqmA structure was built manually using Coot (46) over multiple iterations with further refinement in PHENIX. Images were generated using PyMOL.

### Proteolytic cleavage time course

Apo-VqmA at 5 μM was incubated in the presence of 10 μM synthetic DPO, 10 μM synthetic Ala-AA, or H_2_O, in a final volume of 90 μL for 15 min at RT. A 10 μL aliquot was mixed with stop-solution (2X SDS loading buffer with 5 mM PMSF and 5 mM EDTA) to represent the zero timepoint. Trypsin was added to the preparations at a final concentration of 171 nM and incubated at RT for 1, 2, 3, 5, 10, 15, and 20 min. At each time point, 10 μL of the reactions were added to 10 μL of stop-solution. Samples were heated to 95°C and subjected to SDS-PAGE.

### Thermal shift assay

Thermal-shift analyses of VqmA were performed as described (30). In brief, VqmA was diluted to 5 μM in reaction buffer (20 mM Tris-HCl pH 8, 150 mM NaCl) containing either 10 μM synthetic DPO, 10 μM synthetic Ala-AA, or an equivalent volume of water in an 18 μL reaction (final vol). The mixtures were incubated at RT for 15 min. 2 μL of 200X SYPRO Orange in reaction buffer were added to the samples immediately before analysis. Samples were subjected to a linear heat gradient of 0.05°C s^-1^, from 25°C to 99°C in a Quant Studio 6 Flex System (Applied Biosystems) using the melting-curve setting. Fluorescence was measured using the ROX reporter setting.

### Isothermal Titration Calorimetry (ITC)

A MicroCal PEAQ-ITC (Malvern) instrument was used to measure the binding affinities between VqmA proteins and ligands at 25°C. All synthetic ligands for ITC were prepared in gel filtration buffer (described above). For measurement of Kd, 500 μM ligand was titrated into 50 μM Apo-VqmA or mutant VqmA proteinsusing continuous stirring at 750 rpm. For the ITC-based competition assay, 500 μM synthetic Ala-AA was titrated into 50 μM DPO-VqmA in the presence of 0, 50, 500 μM synthetic DPO, and in the opposite case, 500 μM DPO was titrated into 50 μM Ala-AA-VqmA in the presence of 0, 50, 500 μM Ala-AA. The initial and subsequent 18 injections were 0.4 μL and 2 μL, respectively. Heat changes during the titration were measured by the PEAQ-ITC Control software (MicroCal). Data were fitted and evaluated by the PEAQ-ITC Analysis software (MicroCal).

### Stopped-Flow fluorimetry

A SX20 Stopped Flow Spectrometer (Applied Photophysics) was used for the kinetic study of ligand interactions with VqmA at 25°C. DPO and VqmA have fluorescence spectra of λ_ex_=321 nm/λ_em_=392 nm and λ_ex_=280 nm/λ_em_=340 nm, respectively. For DPO binding to VqmA, the k_obs_ was measured by recording the decrease in DPO fluorescence using a 390 nm emission interference filter. The concentration of VqmA was fixed at 2.5 μM, while DPO was supplied at concentrations from 1-20 μM. For Ala-AA binding to VqmA, the kobs was measured by recording the decrease in VqmA fluorescence using a 302 nm emission interference filter. The concentration of Ala-AA was fixed at 7.5 μM and VqmA, from 3 to 37.5 μM, was supplied. Typically, 5 to 10 traces were recorded and averaged for each data point. Data were fitted with an exponential decay equation using GraphPad Prism6. Formation of complexes follows this equation: k_obs_ = k_on_ x [Analyte] + k_off_. By plotting k_obs_ versus [Analyte], k_on_ and k_off_ are obtained from the slope and Y-axis intercept, respectively.

### Western blot analysis

Western blot analyses probing for FLAG fusions were performed as previously reported (8). *V. cholerae* cells were collected and diluted two-fold in Laemmli sample buffer giving a final concentration of 0.01 OD/μL. Following denaturation for 15 min at 95°C, 0.1 OD_600_ equivalents were subjected to SDS gel electrophoresis. RNAPα was used as loading control. Signals were visualized using an ImageQuant LAS 4000 imager (GE Healthcare).

### Accession code

Atomic coordinates of DPO-VqmA have been deposited in the Protein Data Bank under accession code 6UGL.

## ACKNOWLEDGEMENTS

We would like to thank Dr. Phillip Jeffrey of the Macromolecular Crystallography Core Facility, Dr. Saw Kyin of the Proteomics and Mass Spectrometry Core Facility, and Dr. Venu Vandasvasi of the Biophysics Core Facility, all at Princeton University, for expert assistance. We thank Dr. Frederick Hughson for thoughtful discussions. We thank the Bassler group for insight into the research. The funders had no role in study design, data collection and analysis, decision to publish, or preparation of the manuscript.

## CONFLICT OF INTEREST

The authors declare that they have no conflicts of interest with the contents of this article.

## FOOTNOTES

This work was supported by the Howard Hughes Medical Institute, NIH Grant 5R37GM065859, and National Science Foundation Grant MCB-1713731 (to B.L.B), a Jane Coffin Childs Memorial Fund for Biomedical Research Postdoctoral Fellowship (J.E.P), NIGMS T32GM007388 (O.P.D), a Charlotte Elizabeth Procter Fellowship provided by Princeton University (J.E.S.), and a National Defense Science and Engineering Graduate Fellowship supported by the Department of Defense (J.E.S). This research used resources FMX and AMX of the National Synchrotron Light Source II, a U.S. Department of Energy (DOE) Office of Science User Facility operated for the DOE Office of Science by Brookhaven National Laboratory under Contract No. DE-SC0012704. The Life Science Biomedical Technology Research resource is primarily supported by the National Institute of General Medical Sciences (NIGMS) through a Biomedical Technology Research Resource P41 grant (P41GM111244), and by the DOE Office of Biological and Environmental Research (KP1605010). The authors received expert technical support from Dr. Martin Fuchs and Dr. Jean Jakoncic of NSLS II FMX and AMX beamlines. The content is solely the responsibility of the authors and does not necessarily represent the official views of the National Institutes of Health.

The abbreviations used are: DPO, 3,5-dimethyl-pyrazin-2-ol; Ala-AA, N-alanyl-aminoacetone; QS, quorum sensing; AIs, autoinducers; HCD, high cell density; LCD, low cell density; Tdh, threonine dehydrogenase; AKB, 2-amino-3-ketobutyric acid; ITC, isothermal titration calorimetry; AHLs, acylhomoserine lactones; 3OC_12_-HSL, 3OC_12_-homoserine lactone; EMSAs, electromobility shift assays; d_3_-Ala-AA, (*S*)-2-amino-N-(2-oxopropyl)-3,3,3-d_3_-propanamide; PAS, Per-Arnt-Sim; LBD, ligand-binding domain; HTH, helix-turn-helix motif; DBD, DNA-binding domain; AHP, (2*S*)-2-amino-N-[(2*R*)-2-hydroxypropyl]propanamide; CAI-1, (*S*)-3-hydroxytridecan-4-one; Ea-CAI-1, 3-aminotridec-2-en-4-one; AI-2, 4,5-dihydroxy-2,3-pentanedione; THF, tetrahydrofuran; CDCl_3_, deuterated chloroform; DMSO-d_6_, deuterated dimethylsulfoxide; atm, atmosphere; PE, petroleum ether; EtOAc, ethyl acetate; DCM, dichloromethane; EtOH, ethanol; *t*-Bu, *tert*-butyl; MeOH, methanol; ACN, acetonitrile; Na_2_SO_4_, sodium sulfate; Boc, *tert*-butoxycarbonyl; MTBE, methyl *tert*-butyl ether; Na_2_S_2_O_3_, sodium thiosulfate; HCl, hydrochloric acid; Et_3_N, triethylamine; H_2_, hydrogen gas; Pd(OH)_2_, palladium hydroxide

