## Supplemental Tables1-4 and Figure 1-7 for "Mechanism underlying autoinducer recognition in the *Vibrio cholerae* DPO-VqmA quorum-sensing pathway"

**List of the supporting information:**

**Table S1. Summary of crystal data collection and refinement statistics.**
**Table S2. DALI-based structural homology analysis of VqmA.**
**Table S3. $K_d$ values for Apo-VqmA binding to DPO and analogs.**
**Table S4. Strains used in this study.**
**Table S5. Plasmids used in this study (attached as separate Excel file).**
**Table S6. Primers used in this study (attached as separate Excel file).**
**Figure S1. VqmA does not depend on its cognate ligand to fold and become soluble.**
**Figure S2. Ala-AA binds VqmA without converting into DPO.**
**Figure S3. Ala-AA is not the biological precursor to DPO.**
**Figure S4. The VqmA K185A, K193A, and R195A mutants are unable to bind DNA.**
**Figure S5. AHP does not activate nor bind VqmA.**
**Figure S6. *In vivo* analyses of the activity of mutant VqmA variants.**
**Figure S7. Spectral emission properties of DPO and VqmA.**

**Table S1. Summary of crystal data collection and refinement statistics.**

| Dataset | Native | Au |
|---|---|---|
| Space Group | P $2_12_12_1$ | P $2_12_12_1$ |
| Unit Cell Dimensions (Å) | a=50.4, b=84.6, c=116.8; α=89.9$^{o}$, β=90.0$^{o}$, γ=89.9$^{o}$ | a=50.5, b=84.5, c=114.8; α=89.9$^{o}$, β=90.1$^{o}$, γ=90.0$^{o}$ |
| Wavelength (Å) | 0.920094 | 1.03937 |
| Resolution (Å) | 28.9-2.03 (2.10-2.03) | 30.0-2.15 (2.14-2.13) |
| Unique Reflections | 32520 | 27939 |
| Completeness (%) | 98.67 (89.85) | 97.0 (81.8) |
| <Redundancy> | 8.5 (8.5) | 12.8 (12.2) |
| $R_{meas}$ (%) | 15.8 (78.0) | 10.1 (66.2) |
| < I > / < sigma > | 12.56 (2.64) | 16.42 (3.50) |
| Total atoms | 3225 | |
| $R_{work}$ (%) | 27.6 (37.7) | |
| $R_{free}$ (%) | 33.2 (46.3) | |
| Average B-factor (Å$^2$) | 34.51 | |
| **R.m.s. deviation from ideality** | | |
| Bond lengths (Å) | 0.008 | |
| Bond angles ($^{o}$) | 0.909 | |
| Dihedral angles ($^{o}$) | 3.607 | |
| **Phi-Psi values (Ramachandran)** | | |
| Most favored (%) | 98.22 | |
| Additionally allowed (%) | 1.78 | |
| Outliers (%) | 0 | |

**Table S2. DALI-based structural homology analysis of VqmA.**

| Name (function) | Z-score[a] | Rmsd[b] | Lali[c] | Nres[d] | ID% | PDB Code |
|---|---|---|---|---|---|---|
| HTR-like protein (histidine kinase) | 12.7 | 2 | 95 | 100 | 13 | 3fc7 |
| ThkA (histidine kinase) | 12.1 | 1.9 | 93 | 96 | 17 | 3aos |
| CviR (transcriptional regulator) | 12 | 12.3 | 78 | 258 | 22 | 3qp6 |
| TodS (histidine kinase) | 11.8 | 2.7 | 107 | 128 | 17 | 5hww |
| VraR (transcriptional regulator) | 11.8 | 26.4 | 74 | 208 | 28 | 4gvp |
| PpsR (transcriptional regulator) | 11.7 | 3.1 | 111 | 373 | 14 | 4hh2 |
| Sensor protein (histidine kinase) | 11.6 | 4.4 | 100 | 126 | 15 | 3mxq |
| RcsB (transcriptional regulator) | 11.2 | 8.5 | 66 | 201 | 27 | 5vxn |
| Sensor protein (histidine kinase) | 11.1 | 2.8 | 105 | 116 | 8 | 2r78 |
| Photoactive yellow protein | 11.1 | 2.4 | 103 | 125 | 13 | 1f98 |
| Sensor protein/GGDEF domain | 11.1 | 3 | 111 | 120 | 15 | 3lyx |
| RfpR (sensor protein) | 11.1 | 2.8 | 100 | 103 | 16 | 6dgg |
| VpsT (transcriptional regulator) | 10.7 | 10.2 | 111 | 217 | 23 | 3kln |

[a]Z-score: Measure of similarity between VqmA and target structure

[b]Rmsd: Root mean square deviation of α-carbons between the two structures

[c]Lali: Length of structural alignment in residues

[d]Nres: Total number of amino acids in the hit protein

**Table S3. $K_d$ values for Apo-VqmA binding to DPO and analogs.**

| Ligand | $K_d$ (μM) |
|---|---|
| DPO | 2.3 ± 0.1 |
| 1 | 10.3 ± 3.1 |
| 2 | 7.0 ± 3.1 |
| 3 | 27.8 ± 4.6 |
| 4 | 16.7 ± 5.5 |
| 5 | 6.0 ± 1.0 |
| 6 | 8.3 ± 1.5 |
| 7 | Not detectable |
| 8 | Not detectable |
| 9 | Not detectable |
| 10 | Not detectable |
| 11 | Not detectable |
| 12 | Not detectable |
| 13 | Not detectable |
| 14 | Not detectable |
| 15 | Not detectable |

**Table S4. Strains used in this study.**

| Strain | Genotype | Reference |
|---|---|---|
| *V. cholerae* | Wild-type C6706 | (Thelin and Taylor, 1996) |
| *V. cholerae*, JSS-827 | Wild-type C6706, P*vqmR-lux*::*lacZ* | |
| *V. cholerae*, JSS-826 | C6706, Δ*tdh*, P*vqmR-lux*::*lacZ* | This study |
| *V. cholerae*, JSS-828 | C6706, Δ*vqmA*, P*vqmR-lux*::*lacZ* | This study |
| *V. cholerae*, JSS-852 | C6706, Δ*tdh* Δ*vqmA*, P*vqmR-lux*::*lacZ* | This study |
| *E. coli* BL21(DE3) | *E. coli str. B, F- ompT hsdSB (rBmB-) gal dcm (DE3)* | Agilent |
| *E. coli* BL21(DE3), *tdh*::*kan* | *E. coli str. B, F- ompT hsdSB (rBmB-) gal dcm (DE3) tdh*::*kan* | This study |
| *E. coli* BW25113, Δ*tdh* | *lacIq rrnBT14* Δ*lacZWJ16 hsdR514* Δ*araBADAH33* Δ*rhaBADLD78, tdh*::*kan*$^R$ *(cured)* | (Silpe and Bassler, 2019) |
| *E. coli* TOP10 | *F– mcrA* Δ*(mrr-hsdRMS-mcrBC) φ80lacZ*Δ*M15 ΔlacX74 recA1 araD139* Δ*(ara-leu)7697 galU galK λ– rpsL(Str*$^R$*) endA1 nupG* | Invitrogen |
| *E. coli* S17-1 λ*pir* | D*lacU*169 (Φ*lac*ZDM15) *recA*1 *endA*1 *hsdR*17 *thi*-1, *gyrA*96 *relA*1 λ*pir* | (de Lorenzo and Timmis, 1994) |

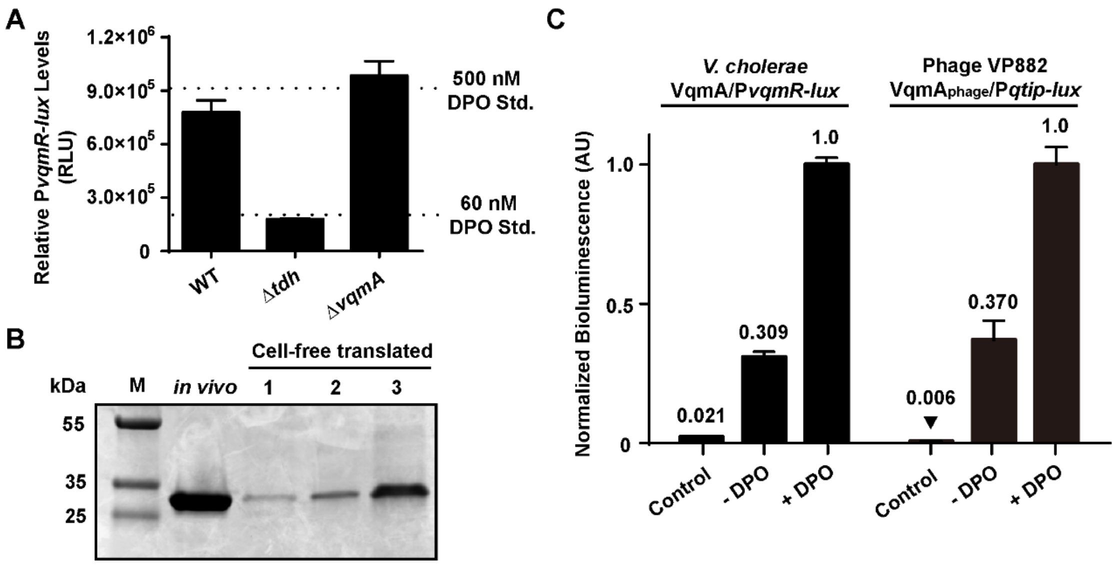


**Figure S1. VqmA does not depend on its cognate ligand to fold and become soluble.** (A) P*vqmR-lux* activity following addition of 25% (v/v) cell-free culture fluids prepared from the indicated *V. cholerae* strains. Dotted lines show the mean reporter activity with n = 3 technical replicates in response to DPO at the designated concentrations. 60 nM DPO was the limit of detection for the reporter strain. (B) SDS-PAGE analysis of His-VqmA produced *in vivo* or by 30, 50, or 90 min (designated 1, 2, 3, respectively) of *in vitro* translation. M = molecular weight marker. (C) Normalized reporter activity for *E. coli* strains harboring the *V. cholerae* VqmA/P*vqmR-lux* or the VP882 phage-encoded $VqmA_{Phage}$/P*qtip-lux* system. Conditions as in Fig. 2. Data in (A) and (C) are represented as mean ± SD with n = 3 biological replicates.

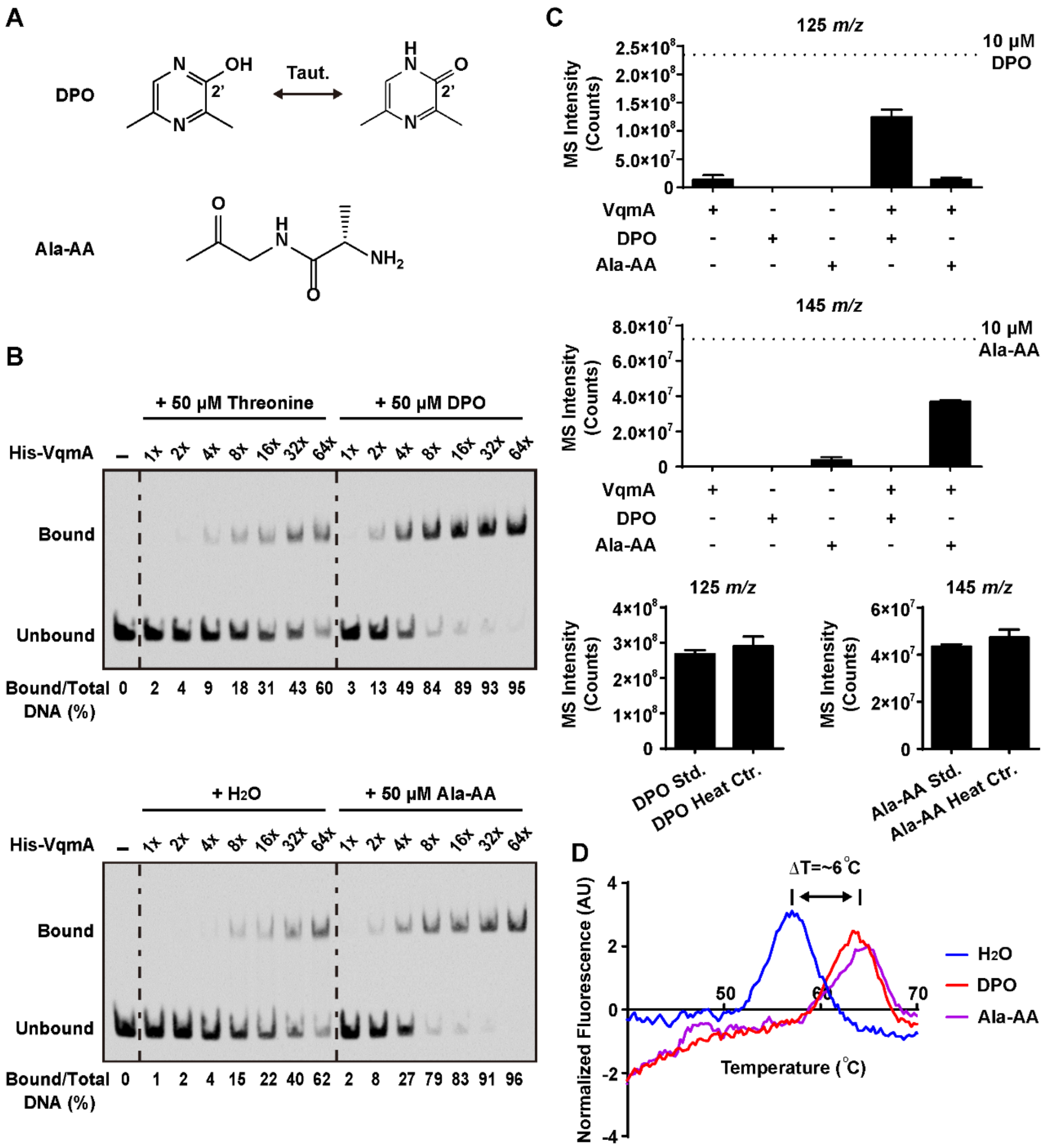


**Figure S2. Ala-AA binds VqmA without converting into DPO.** (A) The structures of DPO and Ala-AA. (B) EMSA analysis of *in vitro* translated His-VqmA binding to P*vqmR* promoter DNA in the presence of the designated ligands or water. Probe and protein concentrations as in Fig. 1 (C). (C) LC-MS analysis of 10 µM DPO (125 *m/z*, top) and 10 µM Ala-AA (145 *m/z*, middle) incubated with or without 50 µM purified Apo-VqmA protein in EMSA buffer, followed by purification of the VqmA protein and release of bound ligand by heating to 70°C. The dotted lines designate the mean with n = 3 technical replicates for 10 µM DPO (top panel) and 10 µM Ala-AA (middle panel) standards prepared in EMSA buffer. When VqmA is omitted (second and third columns in the panels), no or almost no ligand is recovered following the purification steps. Approximately 100 nM DPO is detected in the samples containing 50 µM purified Apo-VqmA (first bar in top panel). The bottom two panels show the results for 10 µM DPO and 10 µM Ala-AA incubated alone in EMSA buffer without and with heating to 70°C. Data are represented as mean ± SD with n = 3 biological replicates. (D) Thermal shift analyses using Apo-VqmA and exogenously supplied $H_2O$ (blue), DPO (red), and Ala-AA (purple). Ligands were provided at 100 μM. Normalized fluorescence represents the first derivative of the raw fluorescence data.

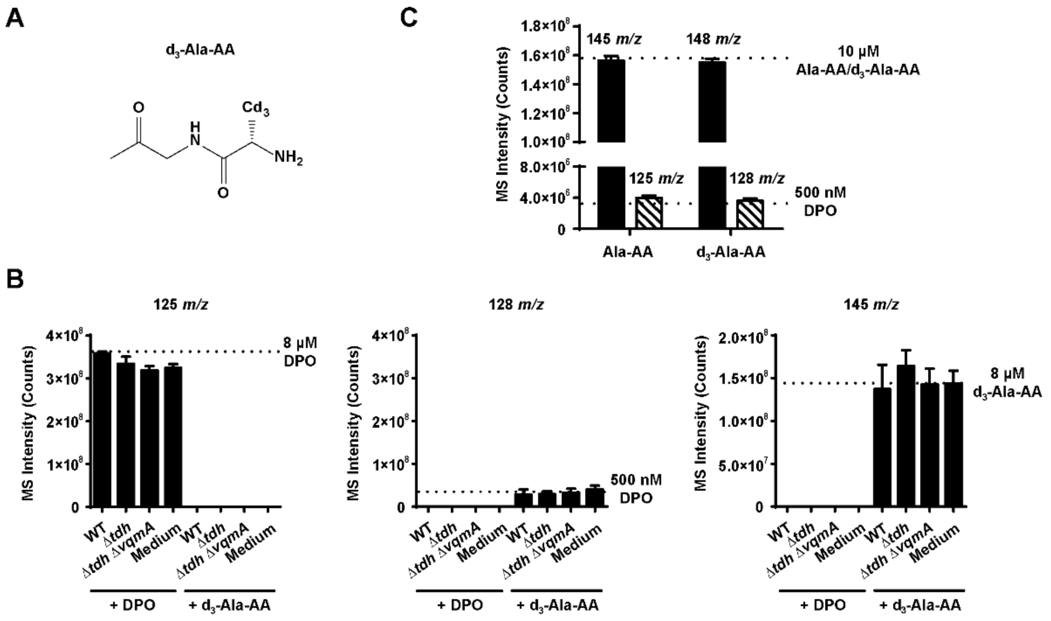


**Figure S3. Ala-AA is not the biological precursor to DPO.** (A) The structure of $d_3$-Ala-AA. (B) LC-MS quantitation of DPO (125 *m/z*, left graph), $d_3$-DPO (128 *m/z*, middle graph), and $d_3$-Ala-AA (148 *m/z*, right graph) extracted from minimal medium or culture fluids from the designated *V. cholerae* strains incubated with 100 µM DPO (left set of bars in each graph) or 100 µM $d_3$-Ala-AA (right set of bars in each graph). Samples were diluted 10-fold prior to analysis. The dotted lines designate the means for DPO and $d_3$-Ala-AA standards at the designated concentrations. (C) LC-MS analysis of 10 µM Ala-AA and 10 µM $d_3$-Ala-AA standards. Quantitation of Ala-AA and $d_3$-Ala-AA (145 *m/z* and 148 *m/z*, black bars), and DPO and $d_3$-DPO (125 *m/z* and 128 *m/z*, striped bars) detected in each standard are shown. Data in (B) are represented as mean ± SD with n = 3 biological replicates and data in (C) are represented as mean ± SD with n = 3 technical replicates.

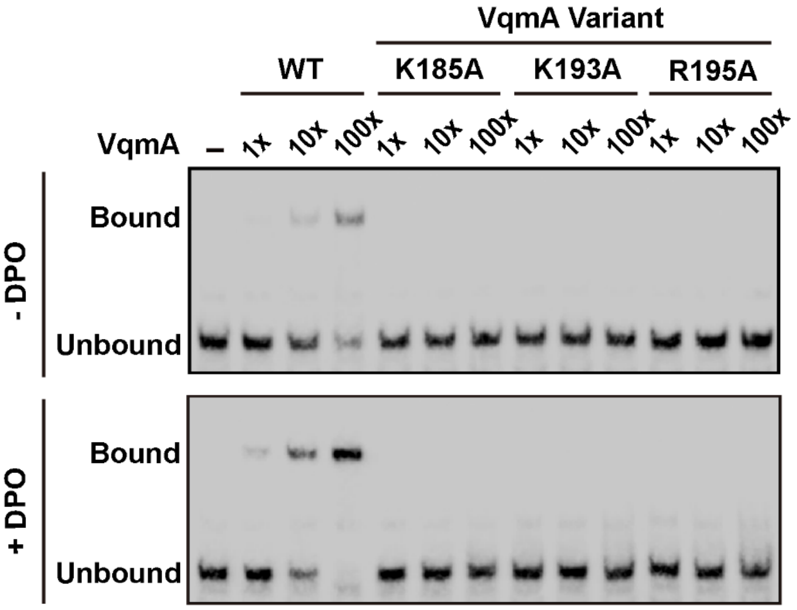


**Figure S4. The VqmA K185A, K193A, and R195A mutants are unable to bind DNA.** EMSA analysis showing P*vqmR* DNA and either WT Apo-VqmA or the indicated Apo-VqmA variants in the absence and presence of DPO (upper and lower panels, respectively). The leftmost lane in each gel shows the no protein control. Relative VqmA concentrations are 5, 50, and 500 nM, which correspond to 1, 10, and 100x, respectively.

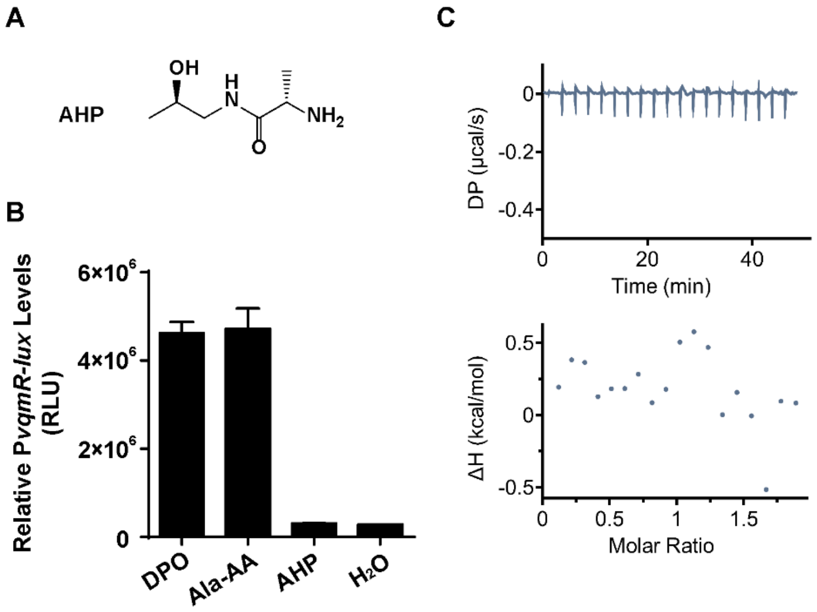


**Figure S5. AHP does not activate nor bind VqmA.** Effect of AHP (A) on VqmA as measured by the P*vqmR* reporter (B) and ITC (C) assays.

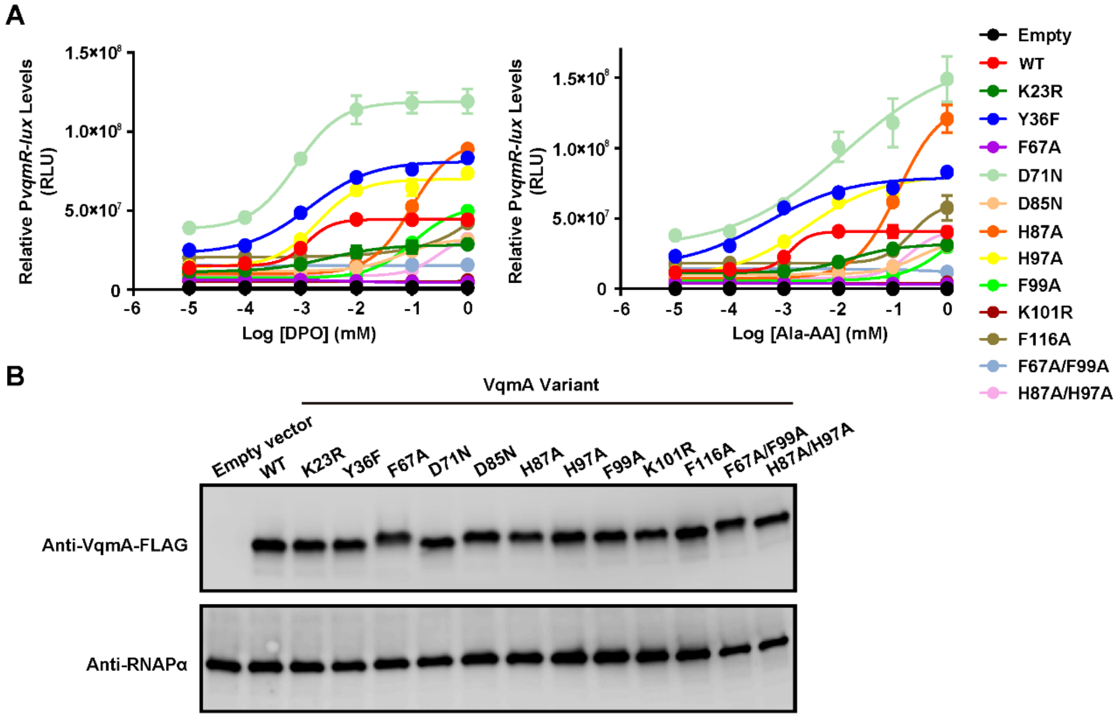


**Figure S6. *In vivo* analyses of the activity of mutant VqmA variants.** (A) Dose response analyses for WT VqmA and mutant VqmA variants using the P*vqmR-lux* reporter strain. DPO (left) and Ala-AA (right) were provided at the specified concentrations. (B) Western blot showing production of the designated VqmA-3xFLAG proteins. The RNAPα subunit was used as the loading control.

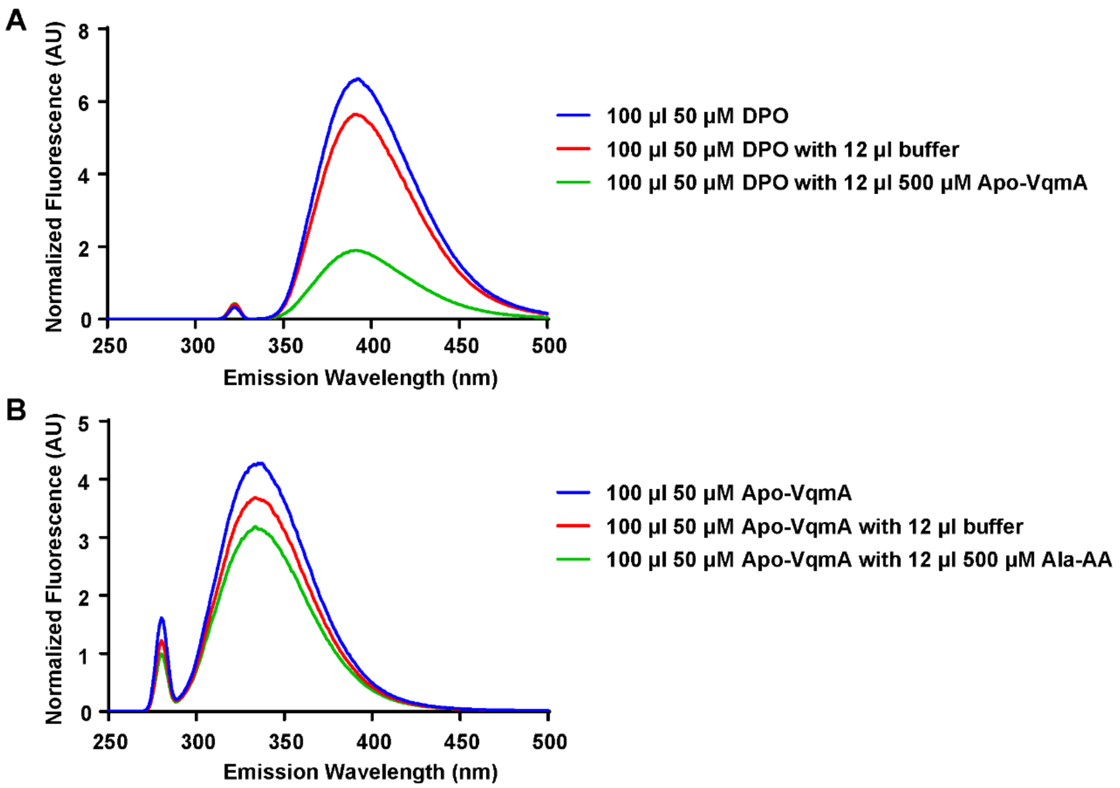


**Figure S7. Spectral emission properties of DPO and VqmA.** (A) When excited at 321 nm, DPO emission, which occurs at 392 nm, decreases upon binding to Apo-VqmA. (B) When excited at 280 nm, Apo-VqmA emission, which occurs at 340 nm, decreases following Ala-AA binding.
